# The effect of linker conformation on performance and stability of a two-domain lytic polysaccharide monooxygenase

**DOI:** 10.1101/2023.06.01.543078

**Authors:** Zarah Forsberg, Anton A. Stepnov, Giulio Tesei, Yong Wang, Edith Buchinger, Sandra K. Kristiansen, Finn L. Aachmann, Lise Arleth, Vincent G. H. Eijsink, Kresten Lindorff-Larsen, Gaston Courtade

**Affiliations:** Faculty of Chemistry, Biotechnology and Food Science, Norwegian University of Life Sciences (NMBU), 1432 Ås, Norway; Structural Biology and NMR Laboratory, Linderstrøm-Lang Centre for Protein Science, Department of Biology, University of Copenhagen, Denmark; College of Life Sciences, Zhejiang University, Hangzhou, 310027, China; Vectron Biosolutions AS, Abels gate 5, 7030 Trondheim, Norway; Department of Biotechnology and Food Science, NTNU Norwegian University of Science and Technology, 7491 Trondheim, Norway; X-ray and Neutron Science, Niels Bohr Institute, University of Copenhagen, Copenhagen, Denmark

**Keywords:** lytic polysaccharide monooxygenase, flexible linkers, carbohydrate-active enzyme, multidomain

## Abstract

A considerable number of lytic polysaccharide monooxygenases (LPMOs) and other carbohydrate-active enzymes are modular, with catalytic domains being tethered to additional domains, such as carbohydrate-binding modules, by flexible linkers. While such linkers may affect the structure, function, and stability of the enzyme, their roles remain largely enigmatic, as do the reasons for natural variation in length and sequence. Here, we have explored linker functionality using the two-domain cellulose-active *Sc*LPMO10C from *Streptomyces coelicolor* as a model system. In addition to investigating the wild-type enzyme, we engineered three linker variants to address the impact of both length and sequence and characterized these using SAXS, NMR, MD simulations, and functional assays. The resulting data revealed that, in the case of *Sc*LPMO10C, linker length is the main determinant of linker conformation and enzyme performance. Both the wild-type and a serine-rich variant, which have the same linker length, demonstrated better performance compared to those with either a shorter or longer linker. A highlight of our findings was the substantial thermostability observed in the serine-rich variant. Importantly, the linker affects thermal unfolding behavior and enzyme stability. In particular, unfolding studies show that the two domains unfold independently when mixed, while the full-length enzyme shows one cooperative unfolding transition, meaning that the impact of linkers in biomass processing enzymes is more complex than mere structural tethering.

## Introduction

Enzymatic depolymerization of polysaccharides in lignocellulosic biomass, in particular cellulose, is of great scientific and industrial interest to produce biofuels, biomaterials and commodity chemicals (Sheldon, 2016). However, the recalcitrance of crystalline cellulose poses an obstacle for its efficient saccharification (Rubin *et al*, 2007). This obstacle may be overcome by applying lignocellulolytic enzyme cocktails comprising glycoside hydrolases (GHs, e.g., xylanases, cellulases) and copper-dependent redox enzymes known as lytic polysaccharide monooxygenases (LPMOs). Whilst GHs hydrolyze their substrates, LPMOs use an oxidative mechanism to cleave the β-1,4 glycosidic bonds (Vaaje-Kolstad *et al*, 2010; Horn *et al*, 2012) in cellulose and various hemicellulosic β-glucans (Agger *et al*, 2014). Many carbohydrate-active enzymes are modular, consisting, for example, of a catalytic and a carbohydrate-binding module (CBM). These domains are connected by linker sequences of variable amino acid composition and length (Gilbert & Hazlewood, 1993; Sammond *et al*, 2012). While there is extensive and in-depth understanding of structure-function relationships for the individual catalytic and carbohydrate-binding modules, knowledge on the conformations and roles of linkers remain scant, despite pioneering work on fungal GH7 cellulases (Payne *et al*, 2013; Kołaczkowski *et al*, 2020).

Multidomain LPMOs with a family AA10 catalytic domain are often tethered by disordered linkers of different lengths to one or more catalytic or substrate-binding domains. For example, two chitin-binding modules (CBM5 and CBM73) in *Cj*LPMO10A (Forsberg *et al*, 2016), a GH18 and a CBM5 in *Jd*LPMO10A (Mekasha *et al*, 2020), two FnIII domains and a CBM5 in *Bt*LPMO10A (Manjeet *et al*, 2019), or a CBM2 in *Sc*LPMO10C as in our model enzyme. It is well established that the CBM2 domain enhances binding affinity of the full-length enzyme to its cellulose substrate (Courtade *et al*, 2018), which is important for LPMOs as localization close to the substrate protects the enzyme from self-inactivating off-pathway reactions (Forsberg & Courtade, 2023). .However, the roles of the linkers in these types of bacterial enzymes remain largely enigmatic.

Intrinsically disordered linkers have heterogenous conformations and characterizing the conformational ensembles of multidomain proteins is crucial to understand their function. Although they may be regarded as just keeping domains close, it is clear that linkers may be key functional regions that govern structurally and functionally important interactions between the folded domains (Sørensen & Kjaergaard, 2019). Linker functionality is affected by the amino acid composition, sequence patterns, and length (Das *et al*, 2015). For example, prolines and charged residues (aspartate, glutamate, arginine and lysine) may promote extended conformations, and glycines promote flexibility and conformational freedom in linkers as well as intrinsically disordered proteins (IDPs) (Das & Pappu, 2013; Sørensen & Kjaergaard, 2019; Marsh & Forman-Kay, 2010). As another example, serine-rich linkers are thought to be more flexible than proline-rich linkers (Shen *et al*, 1991; Howard *et al*, 2004). In the context of LPMOs, Tamburrini *et al* identified that numerous LPMOs have an intrinsically disordered region located at the C-terminus. These C-terminal extensions are typically longer than the linkers found between domains and are unique to LPMOs, not observed in other carbohydrate-active enzymes or oxidoreductases (Tamburrini *et al*, 2021). Most of these LPMO C-terminal extensions feature at least one putative binding site, underpinning the importance of these disordered regions in LPMO function.

Structural characterization of multidomain proteins is challenging because the flexibility of the linker allows for many individual conformations, including different interdomain distances and relative orientations. Therefore, elucidating the structure of multidomain proteins typically requires a combination of complementary biophysical techniques and computer simulations (Thomasen & Lindorff-Larsen, 2022). One key technique in this respect is small-angle X-ray scattering (SAXS), which can reveal the average dimension and conformation of multidomain proteins in solution as well as provide information about the structural dispersion of flexible conformations (Bernadó *et al*, 2007; Karlsen *et al*, 2015). Moreover, SAXS curves can be calculated from atomistic coordinates, allowing integration of experimental SAXS data with computer modelling or simulations (Larsen *et al*, 2020). Molecular dynamics (MD) simulations allow modelling of the experimentally observed data and can yield models with predictive value. Accurate MD simulations hinge on realistic, sufficiently detailed, and computationally expensive models and the use of relevant simulation times, which may lead to prohibitive demands on computational power. In practice, it is desirable to achieve a compromise between accuracy and computational efficiency, which can be attained by using coarse-grained (CG) models. CG models have been successfully used to model IDPs and flexible linkers (Latham & Zhang, 2022).

Here, we use a combination of SAXS and MD simulations, as well as NMR spectroscopy, to investigate the conformation of a two-domain LPMO, namely *Sc*LPMO10C (previously known as CelS2; 34.6 kDa). *Sc*LPMO10C is a cellulose-oxidizing LPMO from *Streptomyces coelicolor* A3(2) (Forsberg *et al*, 2011) that is composed of an N-terminal catalytic domain belonging to family 10 of auxiliary activities in the carbohydrate-active enzyme database (*Sc*AA10; PDB: 4OY7; residues 35–228; 20.9 kDa; (Forsberg *et al*, 2014)), which is connected to a C-terminal family 2 carbohydrate-binding module (*Sc*CBM2; PDB: 6F7E; residues 263– 364; (Courtade *et al*, 2018)) through a linker (34 residues; residues 229–262) that is rich in prolines, threonines and charged residues.

To investigate the effect of linker composition and length we engineered three variants of *Sc*LPMO10C. These variants were designed by substituting the wild-type linker with linkers derived from other carbohydrate-active enzymes. The four linkers are illustrated in Figure 1, where we provide a graphical representation of the domain architecture of *Sc*LPMO10C (AA10-linker-CBM2), and the primary structures of the linkers. Additional information about the linkers is presented in Table 1, and the complete protein sequences including domain boundaries are presented in Table S1. Our engineered variants comprise, a shorter, 20-residue linker of similar amino acid composition as the wild-type linker (shortened linker; SL), an extended, 59-residue linker also of similar composition as the wild-type linker (extended linker; EL), and a serine-rich linker (SR) that has the same number of residues as the WT but is devoid of prolines or charged amino acids. Following the creation of these variants, we used SAXS, NMR, computational modelling, and functional assays to investigate how the structure, dynamics, stability, and catalytic performance of the enzyme is affected by variation in the linker. The results show that linker length may be more important than sequence for determining the conformation and activity of *Sc*LPMO10C. Overall, our findings provide a basis for understanding the structural and functional roles of linkers in not only in LPMOs but also multimodular proteins in general.

**Table 1.**
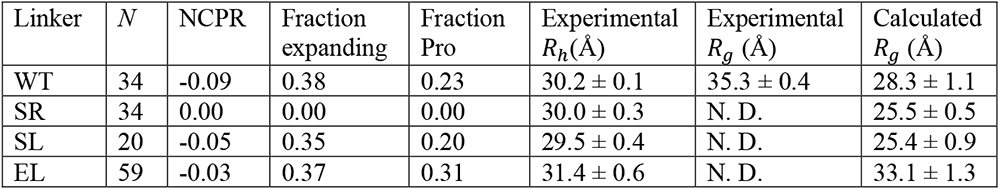
Overview of the properties of four different linker variants of *Sc*LPMO10C. The experimental hydrodynamic radius, *R*_*h*_, was calculated from NMR diffusion experiments. The radius of gyration, *R*_*g*_, was derived from SAXS data and calculated from uniformly weighted CG MD simulations. Errors are indicated as ± S.D. *N*: number of amino acids in the linker region; NCPR: net charge per residue; fraction expanding: fraction of residues which are predicted to contribute to chain expansion (E/D/R/K/P).

| Linker | $N$ | NCPR | Fraction expanding | Fraction Pro | Experimental $R_h$ (Å) | Experimental $R_g$ (Å) | Calculated $R_g$ (Å) |
| --- | --- | --- | --- | --- | --- | --- | --- |
| WT | 34 | -0.09 | 0.38 | 0.23 | $30.2 \pm 0.1$ | $35.3 \pm 0.4$ | $28.3 \pm 1.1$ |
| SR | 34 | 0.00 | 0.00 | 0.00 | $30.0 \pm 0.3$ | N. D. | $25.5 \pm 0.5$ |
| SL | 20 | -0.05 | 0.35 | 0.20 | $29.5 \pm 0.4$ | N. D. | $25.4 \pm 0.9$ |
| EL | 59 | -0.03 | 0.37 | 0.31 | $31.4 \pm 0.6$ | N. D. | $33.1 \pm 1.3$ |

**Figure 1.**
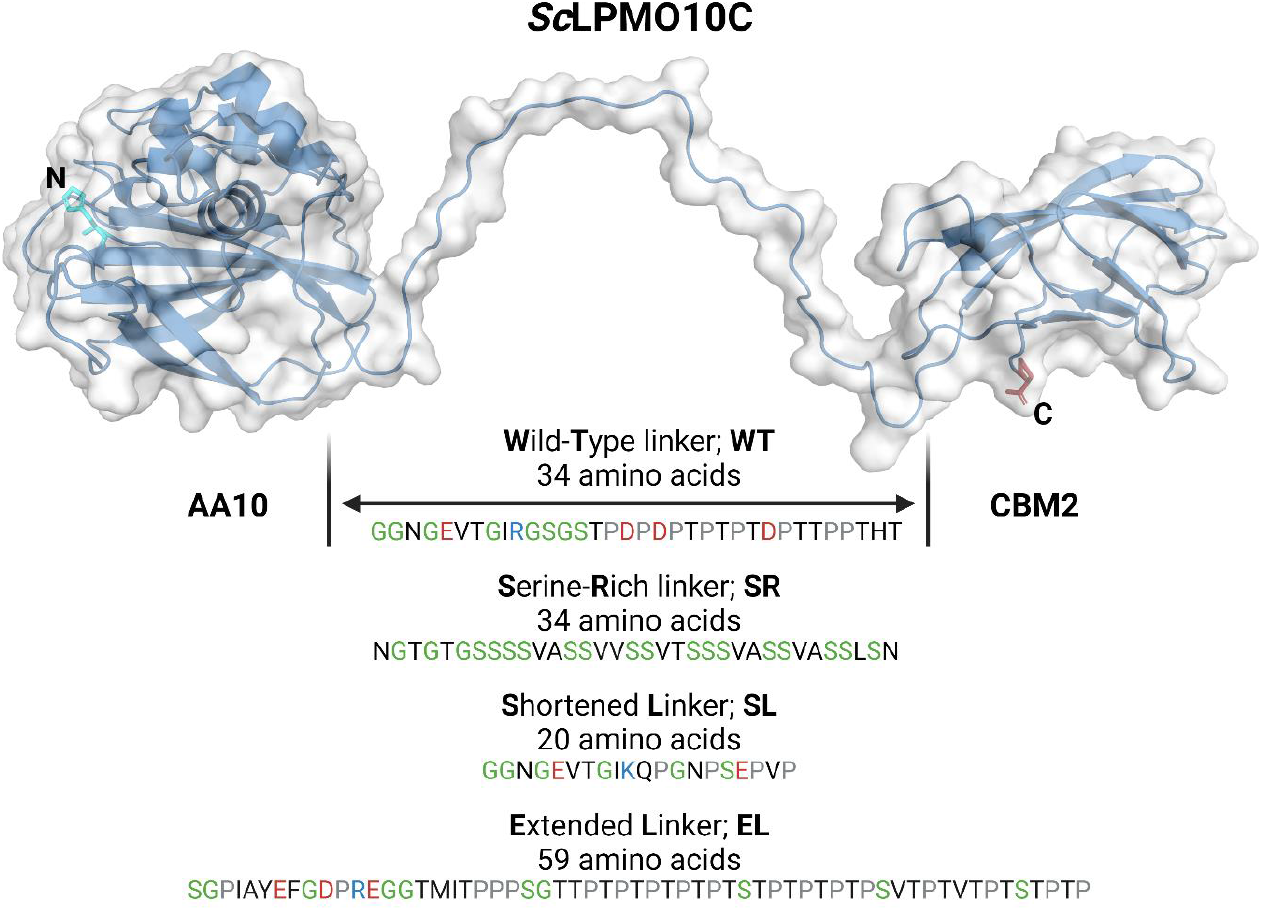
Illustration of the domain architecture of *Sc*LPMO10C with primary structures of the wild-type linker and the three engineered variants. In the linkers, negatively and positively charged residues, as well as prolines, are colored red, blue, and grey, respectively. Glycines and serines are colored green.

## Results

### NMR insights on the linker region of ScLPMO10C

Previously, we published the partial backbone assignment of *Sc*LPMO10C (Courtade *et al*, 2017a) as well as preliminary *T*_1_ and *T*_2_ relaxation, and ^1^H-^15^N NOE data (Courtade *et al*, 2018). In our previous work, approximately 80% of the linker resonances were assigned due to difficulties arising from low signal dispersion of linker resonances combined with signal overlap and a high content of prolines. To enable more detailed studies of the linker region, we have here accomplished > 97 % assignment coverage of the H^N^, N, C^α^, C’ and C^β^ resonances in the linker region. To achieve a high assignment percentage of proline-rich regions in the linker, we used a ^13^C-detected CON-based approach (Jiménez *et al*, 2018) that correlates the amide N atom of an amino acid, *i*, with the carbonyl C’ atom of the preceding amino acid, *i*-1. Using the peak intensities from the *i*-1 residue with respect to Pro, we observed that two prolines in the linker (P248 and P258) had a fraction of cis-Pro in the range of 6–7% (Fig. S1). This is consistent with the fraction of cis-Pro in the IDP α-synuclein (Alderson *et al*, 2018). The updated chemical shift assignment of full-length *Sc*LPMO10C has been deposited in the Biological Magnetic Resonance Data Bank (BMRB) under accession number 27078.

The “average” secondary structure propensity for the linker region was estimated from the chemical shift assignment using TALOS-N, indicating an extended structure with dihedral angles typical of β-strands or polyproline II structures (Fig. 2A), in agreement with previous observations (Courtade *et al*, 2018).

**Figure 2.**
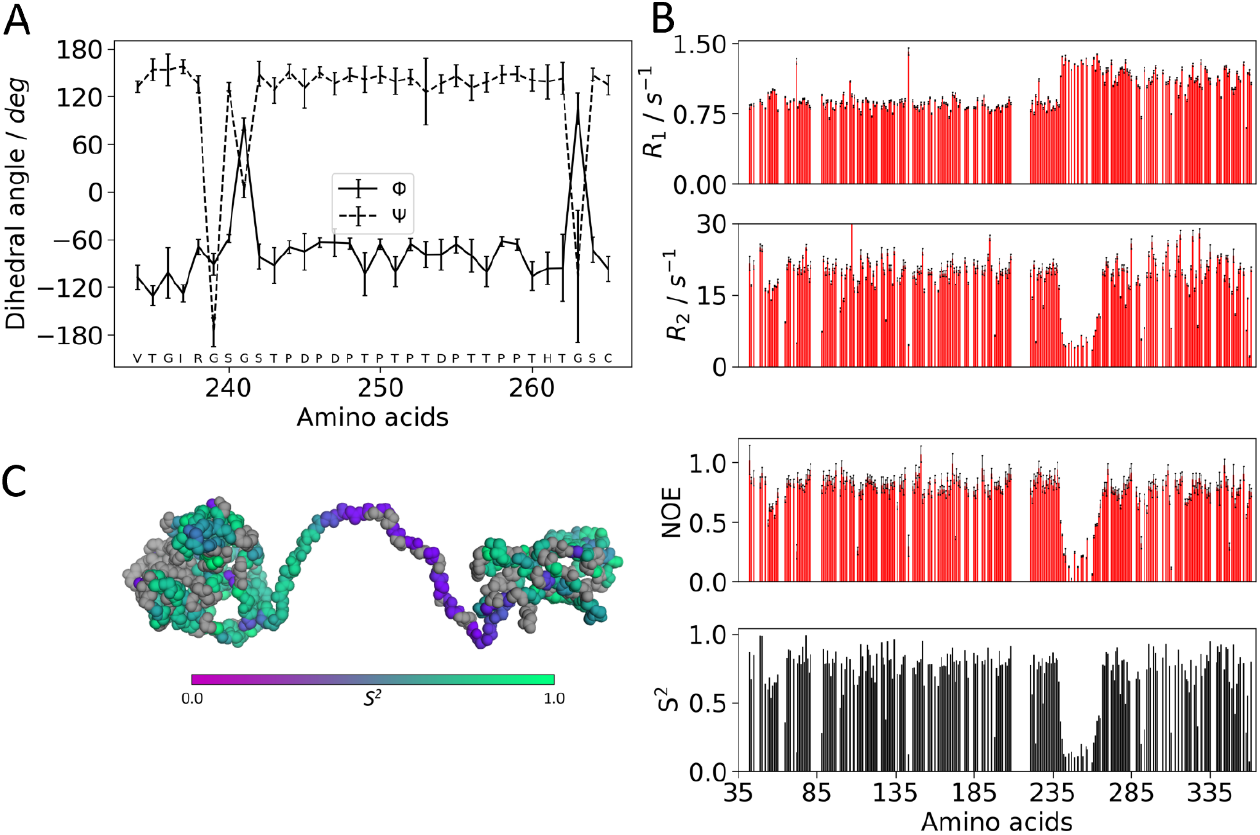
Dihedral angles and ^15^N-relaxation measurements for the *Sc*LPMO10C. (A) Dihedral angles (\phi and \psi) in the linker region and flanking residues of *Sc*LPMO10C derived from the chemical shift assignment using TALOS-N. The values indicate that the linker is in an extended conformation. (B) From top to bottom: ^15^N-R_1_ rates, ^15^N-R_2_ rates, heteronuclear {^1^H}-^15^N NOEs, and calculated generalized order parameters, S^2^. Amino acids displaying flexibility in the ps – ns timescale show increased R_1_, and decreased R_2_, heteronuclear NOEs and S^2^. (C) S^2^ values colored on the structure of *Sc*LPMO10C. Amino acids for which an S2 value could not be determined are colored grey.

Additionally, we analyzed ^15^N relaxation data (*R*_1_, *R*_2_ and ^1^H-^15^N NOE) to provide more complete and accurate insights into the dynamic features of *Sc*LPMO10C, and in particular its newly assigned linker region. Figures 2B and 2C show that amino acids in the linker stand out but display clear features of flexibility in the ps – ns timescale, as can be seen from increased *R*_1_ relaxation rates and decreases in *R*_2_, ^1^H-^15^N NOEs and generalized order parameter, *S*^2^. We note that it is difficult to interpret the *S*^2^ values as internal and overall motions cannot easily be separated for the linker residues.

### Overall dimensions and shape of ScLPMO10C

We studied full-length *Sc*LPMO10C (Fig. 3A–D) and its isolated catalytic domain (called *Sc*AA10; Fig. 3E–F) by SAXS to determine the effect of the linker on the overall conformation and shape of the LPMO. The average sizes of the proteins were determined by calculating the radius of gyration, *R*_*g*_, from experimental SAXS profiles, using Guinier plots. Based on X-ray crystallography (Forsberg *et al*, 2014), SAXS data (Fig. 3E–G), and NMR data showing that *Sc*AA10 is rigid in solution (Fig. 2B), it can be assumed to have a roughly spherical shape. Therefore, the Guinier approximation *qR*_*g*_ ≤ 1.3 (Putnam *et al*, 2007) was used to determine an *R*_*g*_ = 16.25 ± 0.02 Å (Fig. S2). On the other hand, *Sc*LPMO10C is not spherical, and the flexible linker leads to an ensemble of conformations (Courtade *et al*, 2018). Thus, the Guinier approximation was considered to be valid only up to *qR*_*g*_ ≤ 1, an assumption previously reported to be valid for Cel45 from *Humicola insolens*, another two-domain carbohydrate-active enzyme (Receveur *et al*, 2002). By considering this, the *R*_*g*_ of full-length *Sc*LPMO10C was determined to be 35.3 ± 0.4 Å (Table 1; Fig. S2).

**Figure 3.**
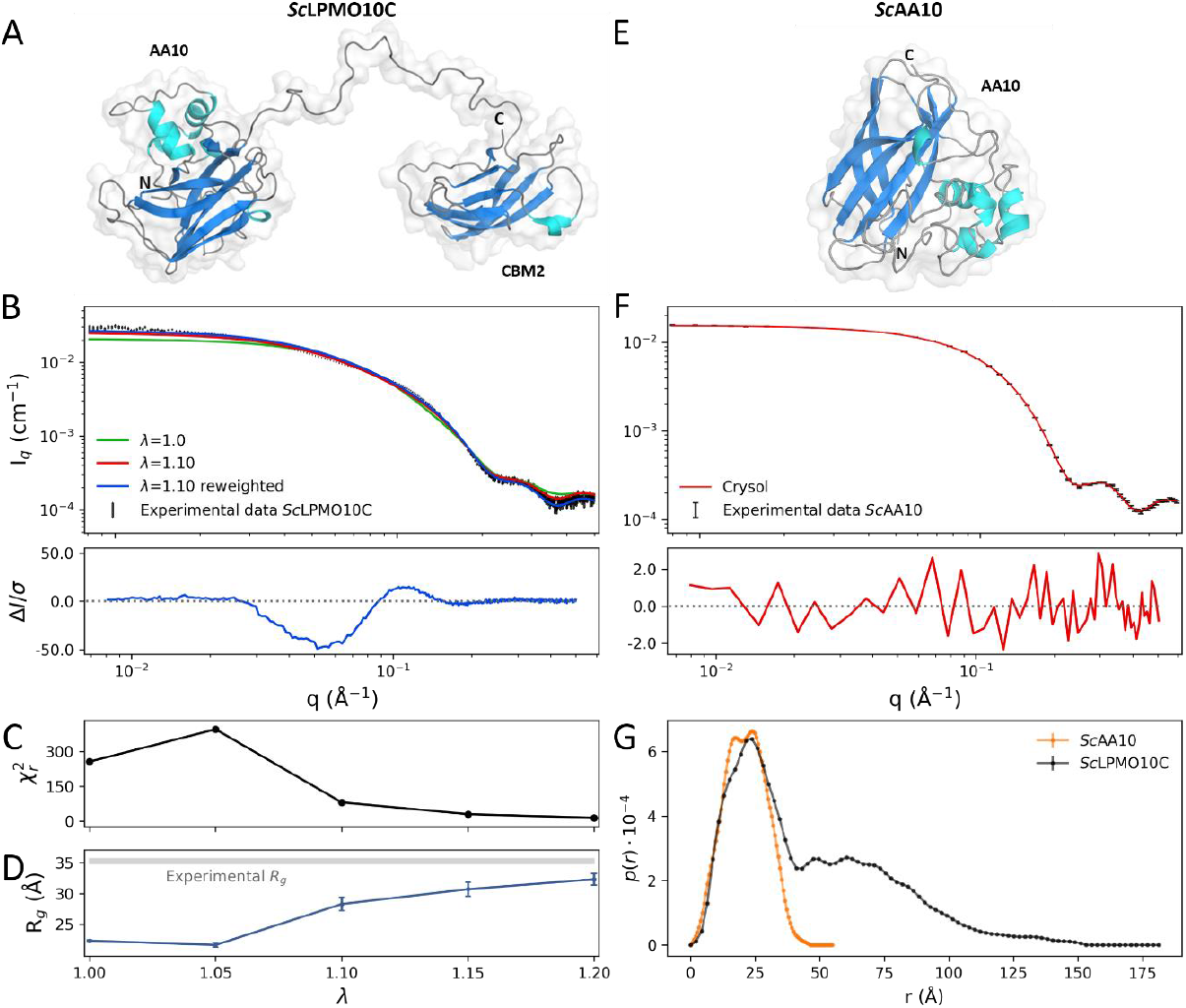
Analysis of the fit to experimental SAXS data of wild-type *Sc*LPMO10C and *Sc*AA10. (A) Representative conformation of *Sc*LPMO10C; the AA10 and CBM2 domains, and the N- and C-termini are labeled. (B) Experimental SAXS data for *Sc*LPMO10C (black), and calculated SAXS data from the ensemble with *λ* = 1.0 (i.e., unmodified protein-water interactions; green), with *λ* = 1.10 (red), and the reweighted ensemble with *λ* = 1.10 (blue). (C) Chi-squared 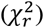 values of the fit to experimental SAXS data of ensembles simulated at different values of *λ*. (D) Calculated radius of gyration (*R*_*g*_) of ensembles simulated at different values of *λ*; the experimentally determined value is indicated by the grey horizontal line. (E) Representative conformation of the isolated catalytic domain, *Sc*AA10 (PDB: 4OY7). (F) Experimental SAXS data for *Sc*AA10 (black), and calculated SAXS data by crysol (red). (G) Distance distribution functions for *Sc*AA10 and *Sc*LPMO10C. Residual plots are shown below panels B and F, where Δ*I* = *I*_*exp*_ *− I*_*fit*_ and σ is the experimental standard deviation. The difference between the experimental and calculated values, even when the simulations fit the data well, is because the experimental value represents the *R*_*g*_ of both the protein and the solvation layer, whereas the calculated value is of the protein only. The molecular mass of the *Sc*LPMO10C and *Sc*AA10 was estimated from the *I*(0) values to 34.6 kDa and 21.4 kDa, respectively.

The distance distribution functions, p(r), for *Sc*AA10 and *Sc*LPMO10C were calculated from their respective scattering intensities. The p(r) profile of *Sc*AA10 exhibits two peaks at respectively 17 Å and 24 Å, and a maximum distance, *D*_*max*_ around 45 Å (Fig. 3G). The p(r) profile of *Sc*LPMO10C displays a biphasic pattern with an initial peak from zero to ∼50 Å, a pronounced shoulder from ∼50 to ∼100 Å, and then a tail that flattens out to a *D*_*max*_ around 150 Å (Fig. 3G). This is consistent with the full-length enzyme having an elongated shape due to the dumbbell shape and the interconnecting flexible linker. The first peak, with a maximum at around *r* = 25 Å represents intramolecular distances within the *Sc*AA10 and *Sc*CBM2 domains. The second part of the curve with the pronounced shoulder stems from (larger) interdomain distances between the *Sc*AA10 and *Sc*CBM2 domains, indicating an extended conformation of the protein (Fig. 3G). The SAXS profile calculated on the structure of *Sc*AA10 (PDB: 4OY7) using CRYSOL displays excellent agreement 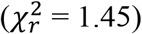 with the experimental data (Fig. 3E)

Despite our efforts, SAXS data for the three linker variants (SR, SL and EL) was not obtained, so their size was instead determined by estimating the hydrodynamic radius, *R*_*h*_, (Table 1) from the translational diffusion coefficient, *D*_*t*_, measured by pulsed-field gradient NMR diffusion experiments (Fig. S3). All proteins have *R*_*h*_ values in the range 29 – 32 Å, and the uncertainties in the measurements are such that there is no discernible difference between the four enzyme variants. For the wild-type, *R*_*g*_/*R*_*h*_ ≈ 1.2, which is consistent with ratios found for expanded conformations in IDPs (Pesce *et al*, 2023; Nygaard *et al*, 2017). This is further evidence that the linker in *Sc*LPMO10C is predominantly extended.

### Coarse-grained MD simulations describe SAXS data

To gain more insight into the conformation of the linkers in the *Sc*LPMO10C variants, and to further investigate potential linker-dependent differences in the overall size of the protein, conformational ensembles of the wild-type and the three linker variants were generated using coarse-grained MD simulations with the Martini version 3.0.beta.4.17 force field (Fig. S4). We used the MD trajectories to calculate ensemble-averaged SAXS curves (Fig. 3B), where the contribution (weight) of each frame to the scattering profile was optimized iteratively using Bayesian-Maximum Entropy (BME) (Pesce & Lindorff-Larsen, 2021) approach. Simulations of the wild-type enzyme resulted in a lower *R*_*g*_value (22 Å; Fig. 3C) compared to the experimentally determined *R*_*g*_ value (35.3 ± 0.4 Å). Consequently, agreement between the experimentally determined and calculated SAXS profiles for the wild type was poor (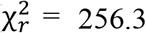; Fig. 3D). Such unphysical compactness in MD simulations using the Martini force field has been observed previously (Larsen *et al*, 2020; Thomasen *et al*, 2022), and is attributed to protein “stickiness” that is a consequence of the force field that promotes too strong protein-protein interactions.

To improve agreement between SAXS data and simulations, we gradually increased the interaction strength between protein beads and water beads (Fig. 3D), as described in the Materials and Methods. Using this approach, *λ* = 1.10 (i.e., a 10% increase of the protein-water interaction strength) resulted in a slight improvement of the fit (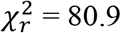; Fig. 3) between the experimental and computed SAXS curves and resulted in a calculated *R*_*g*_= 28.3 ± 1.1 Å (Table 1). Although further incrementations of the protein-water interaction strength (*λ* = 1.15 and *λ* = 1.20) resulted in better agreement with SAXS data, we chose to continue with *λ* = 1.10 as this was a compromise between improving the model and modifying the original force field as little as possible.

We used the simulations to interpret the experimental data by reweighting the trajectory (obtained with *λ* = 1.10) using a BME protocol (Pesce & Lindorff-Larsen, 2021). In this approach, the weights of the simulation are optimized to fit the SAXS experiments. This reweighted ensemble had a calculated *R*_*g*_= 32.3 ± 1.1 Å, and the SAXS curve calculated from the reweighted ensemble had a modest improvement of the fit 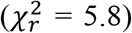 compared to the uniformly weighted ensemble. We note here that the difference between the experimental value (obtained from a Guinier fit to the SAXS experiments) and the calculated values is because the former includes a contribution from the solvent layer, whereas the latter represents the *R*_*g*_ value of the protein only.

Based on these results, we generated uniformly weighted ensembles of the three linker variants by running simulations using *λ* = 1.10. The results (Table 1; Fig. S5) show that EL has the largest *R*_*g*_ value and the most extended conformations, whereas SR and SL have similar *R*_*g*_ values, of around 25 Å, and more compact conformations than EL and WT (Table 1). These results were unexpected because SR (*N* =34) is 14 amino acids longer than SL (*N* = 20), and because both the wild type and SR have linkers of equal length. The results suggest that differences in the calculated *R*_*g*_ values are not only caused by differences in the linker length (number of amino acids), but also by variation in the sequence composition of the linker. This is also supported by the lack of variance in *R*_*h*_ values (Table 1). To gain a better understanding of how linker sequences affect linker length, we conducted an *in silico* analysis to model a large number of linker sequences using recently developed techniques for the analysis of IDPs. The results of this analysis are presented below.

### Sequence length and not composition determines the size of isolated linkers

To obtain a larger coverage in terms of linker lengths and sequence compositions, we searched the UniProtKB Reference Proteomes (Bateman *et al*, 2015) database for linker sequences in proteins with the same domain architecture as *Sc*LPMO10C, i.e. AA10-linker-CBM2. In total, we found 164 unique linkers (including the four discussed above and depicted in Table S1), that we analyzed with MD simulations to investigate whether the *R*_*g*_ of the linker is influenced by its amino acid composition. In these simulations, linkers can be described as random polymer chains (Zheng *et al*, 2020) with R_g_ × k*N*^v^, where *N* is the number of amino acids, k is a constant, and ν is a scaling exponent that is sequence dependent (e.g., ν = ∼0.5 for glycine-serine linkers, ν = ∼0.57 for proline-rich linkers, and ν = ∼0.7 for linkers with charged residues (Sørensen & Kjaergaard, 2019)). Briefly explained, ν = 0.5 is the situation when protein-protein and protein-water interactions balance each other; when protein-water interactions are stronger ν > 0.5, and when protein-protein interactions are stronger, the protein compacts and ν < 0.5. Therefore, if the amino acid composition of the linkers influences the *R*_*g*_ we would not expect a model where all the linkers are described by a single ν to capture the *R*_*g*_ data.

We simulated the 164 linker sequences (average *N* = 40, median *N* = 38, min *N* = 22, max *N* = 72, SD = 11) using CALVADOS, a single-bead model trained with experimental data of IDPs (Tesei *et al*, 2021). From the simulation trajectories of all the sequences, we calculated the *R*_*g*_ in three steps (see Materials and Methods for details). First, the persistence length, *l*_*p*_, (Fig. 4A), which reflects the bending stiffness of the linkers, was calculated by fitting the autocorrelation function of bond vectors to a single exponential decay. Second, these *l*_*p*_ values were used as fixed parameters in the fit of the intramolecular pairwise distances to estimate the scaling exponent, ν (Fig. 4B). We found that the distributions of *l*_*p*_ and ν are both narrow (*l*_*p*_= 0.58 ± 0.04 nm, ν = 0.534 ± 0.009), which means that all sequences are approximately equally disordered and flexible. The calculated value for the scaling exponent is in agreement with previous observations for IDPs (ν = 0.51–0.60) (Marsh & Forman-Kay, 2010). Third, the resulting *R*_*g*_ data calculated solely on the average scaling exponent, i.e., *R*_*g*_ = 0.23 × *N*^0.534^, was found to accurately capture the dependence of *R*_*g*_ on sequence length (Fig. 4C–D). This means that for these 164 sequences the amino acid composition does not influence linker conformation.

**Figure 4.**
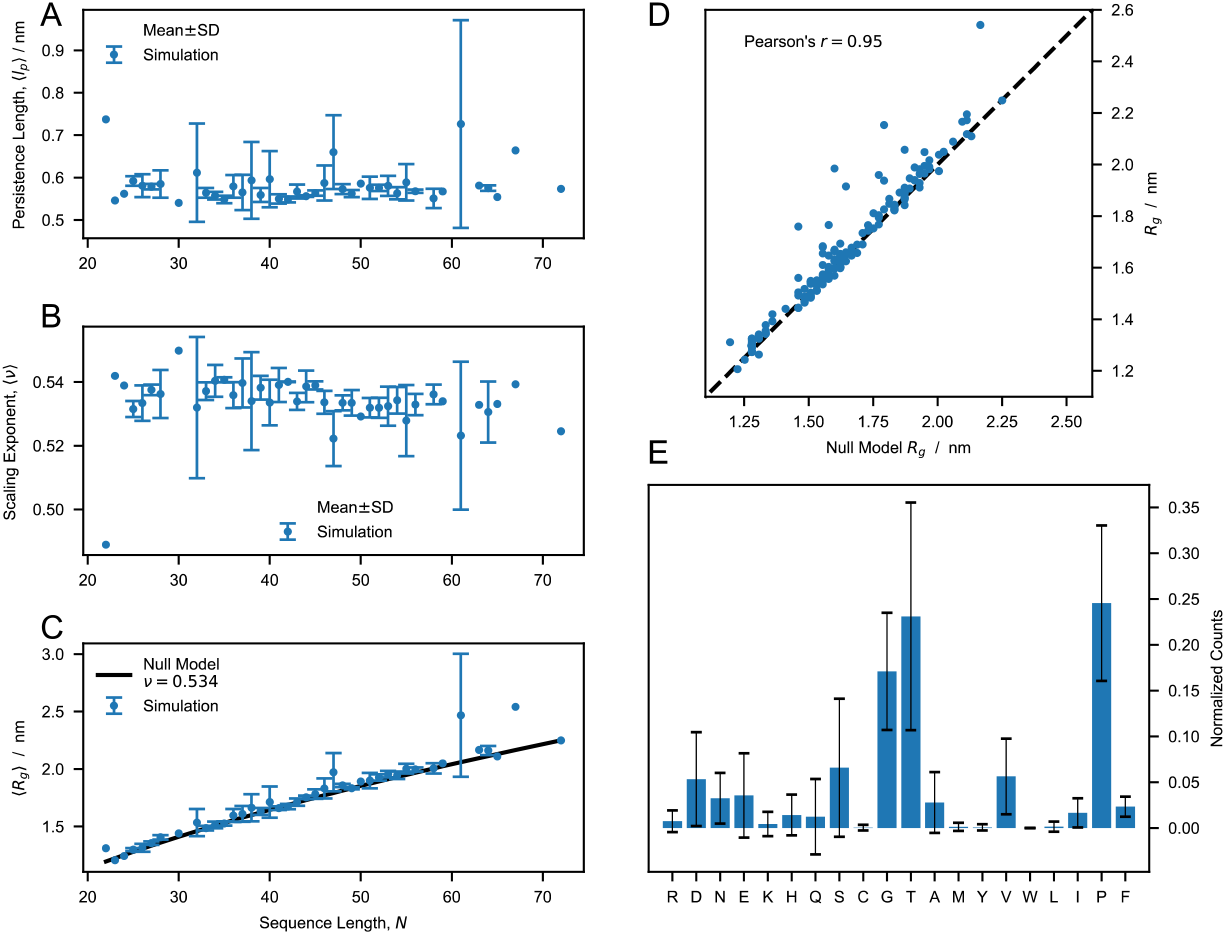
Analysis of conformational ensembles of 164 LPMO linkers. (A and B) Average persistence length, *l*_*p*_, (A) and scaling exponent, ν, (B) for linkers of equal sequence length, *N*. Error bars are SD, meaning that the graphs show mean ± SD. (C) Comparison between the average *R*_*g*_for linkers of *N* and prediction of the null model, i.e. *R*_*g*_ = 0.23 × *N*^0.534^ nm. (D) Correlation between the *R*_*g*_values of the 164 linkers and the corresponding predictions of the null model. (E) Overall amino acid composition of the 164 sequences. The error bars show the SD of the distributions calculated for single linker sequences.

### The linker region affects enzyme stability under turnover conditions

To probe the functionality of the linker variants, cellulose solubilization reactions were carried out in the absence or presence of exogenously added hydrogen peroxide (H_2_O_2_). In the experiment with no H_2_O_2_ added (Fig. 5A), all LPMOs displayed similar substrate solubilization rates during the first 24 hours of incubation. However, after 24 hours, product release by SL and EL slowed down significantly compared to the other two enzymes, which is indicative of LPMO inactivation. The introduction of hydrogen peroxide to LPMO reactions led to much (at least 10–100-fold) faster substrate oxidation rates (Fig. 5B), as expected (Bissaro *et al*, 2017; Chang *et al*, 2022; Kont *et al*, 2020). Interestingly, while the two variants with equally long linkers, WT and SR, both converted essentially all added H_2_O_2_ to oxidized products at similar speeds (i.e., about 100 μM of oxidized products after 10 minutes), SL and EL were much less effective. The latter is indicative of off pathway reactions leading to enzyme inactivation that are promoted when an LPMO is exposed to H_2_O_2_ while not optimally interacting with the substrate (Loose *et al*, 2018; Courtade *et al*, 2018; Stepnov *et al*, 2022b). The two LPMO variants with linkers of equal length, but very different sequences, are functionally highly similar.

**Figure 5.**
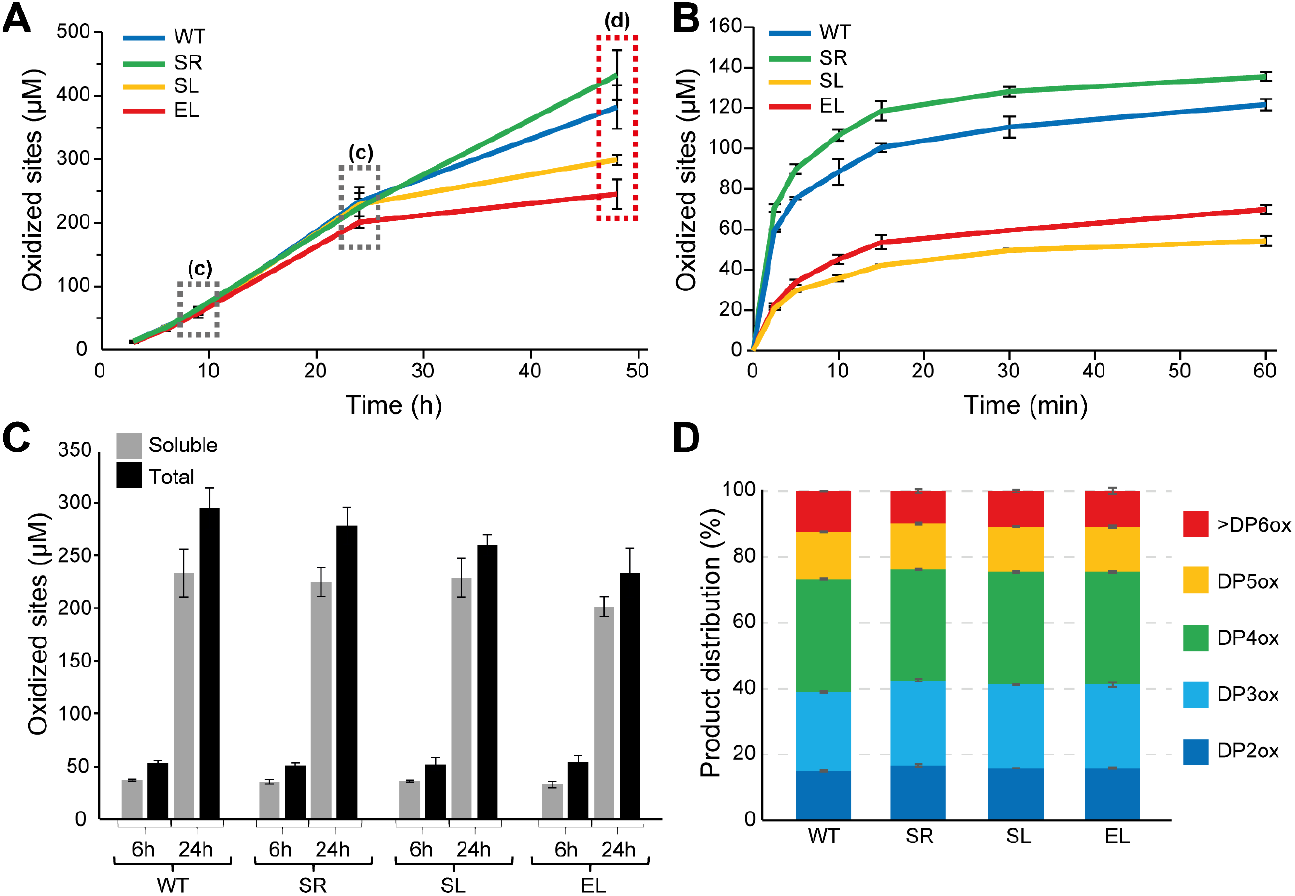
Catalytic activity of four LPMO variants in the absence or presence of exogenously added H_2_O_2_. (A) Progress curves for the formation of soluble oxidized products using 1 mM gallic acid to fuel the reactions. (B) Similar reaction as in panel A, but with exogenous H_2_O_2_ (100 μM) added together with 1 mM ascorbic acid to initiate the reactions. Note the very different time scales in panels A and B. (C) Soluble oxidized products (grey bars) and total amount of oxidized products (soluble + insoluble fraction; black bars), determined for the reactions depicted in panel A at the time-points labelled with dashed boxes in panel A. (D) Degree of polymerization of the soluble oxidized products (DPox) generated after 48 h in the reactions depicted in panel A. All reactions were carried out with 0.5 μM LPMO and 10 g/L Avicel in 50 mM sodium phosphate buffer, pH 6.0. The error bars show ±S.D. (n=3).

Earlier studies with wild-type *Sc*LPMO10C and its CBM- and linker-free truncated variant have shown that, with the substrate concentrations used here, the presence of the CBM promotes localized substrate oxidation, leading to a higher fraction of soluble oxidized products (relative to insoluble products) that on average are shorter (Courtade *et al*, 2018). Figures 5C and 5D show that there are no substantial differences between the four variants in terms of the fraction of soluble oxidized products nor the degree of polymerization of these products.

### The linker region affects thermal unfolding and thermostability

Another possible functional role of the linker relates to structural stability which could be affected by linker-mediated interdomain interactions. Therefore, we used differential scanning calorimetry (DSC) to investigate the conformational stability of *Sc*LPMO10C and its isolated domains, i.e., *Sc*AA10 and *Sc*CBM2. Fig. 6 shows that the apparent melting temperature, *T*_*m*(app)_, of WT (*T*_*m*(app)_ = 65.8 °C) is higher than for *Sc*AA10 (*T*_*m*(app)_ = 61.8 °C) and *Sc*CBM2 (*T*_*m*(app)_ = 52.4 °C). Importantly, the full-length enzyme showed a single transition (Figs. 4A, B), whereas DSC scans of a mixture of *Sc*AA10 and *Sc*CBM2 (Fig. 6C), showed a double transition, at lower temperature compared to the full-length enzyme, with the signal amounting to the sum of the signals obtained for the individual domains. Together, these results clearly show that the linker affects structural stability and mediates domain interactions in the full-length enzyme.

**Figure 6.**
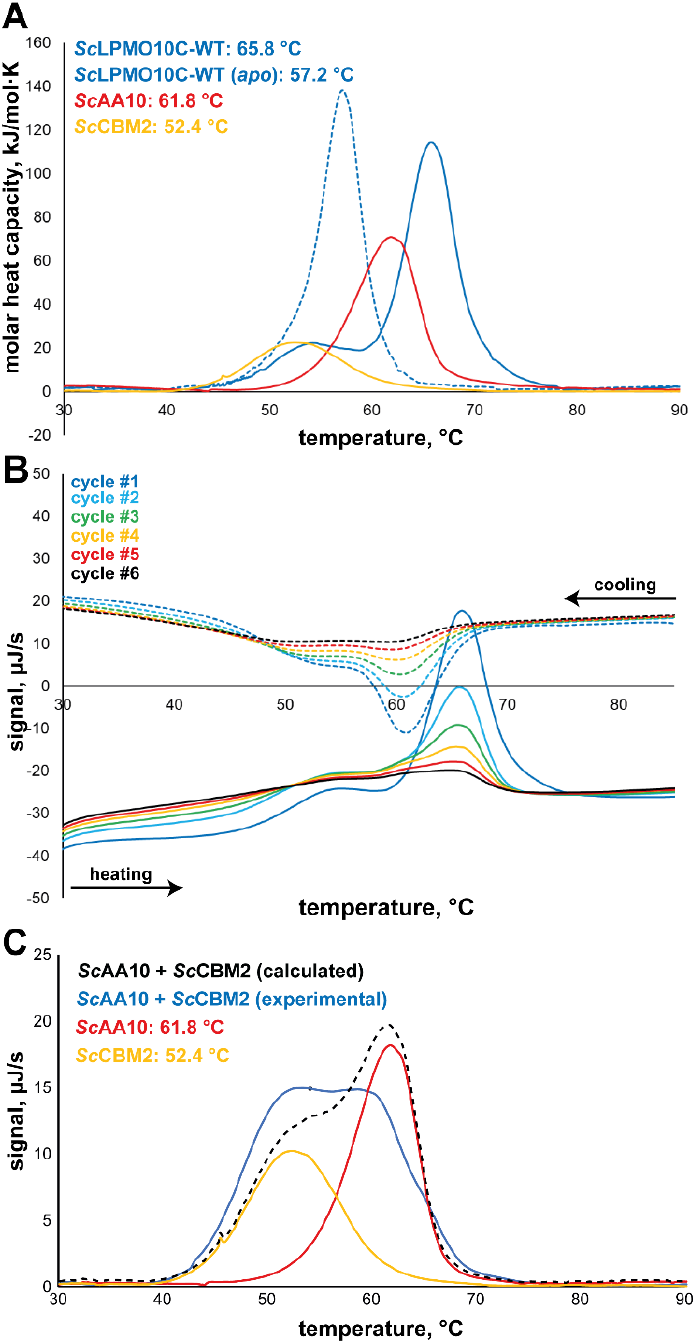
DSC thermograms showing unfolding of *Sc*LPMO10C. (A) Thermal unfolding for the wild-type enzyme, the catalytic domain (*Sc*AA10) and *Sc*CBM2, and their respective calculated apparent *T*_*m*_ values. The molar heat capacity of the protein samples after buffer baseline subtraction is plotted against the temperature. A melting curve of *apo-*enzyme (i.e., copper free) was also recorded (dashed thermogram) showing an expected decrease in apparent *T*_*m*_. The shoulder in the curve for *holo*-*Sc*LPMO10C likely represents a small fraction of *apo*-enzyme. (B) Multiple unfolding and refolding cycles for *Sc*LPMO10C showing some degree of reversibility and a consistent apparent *T*_*m*_. Fig. S6 shows similar data for *Sc*AA10 and *Sc*CBM2. (C) Thermal unfolding of the individual domains, alone, or when combined; the dashed curve shows an unfolding curve calculated by summing the curves obtained for the individual domains. Panel B and C show raw, unprocessed data in which buffer baseline was not subtracted from the signal. All experiments were conducted in sodium phosphate buffer, pH 6.0, at a protein concentration of 1 g/L (*Sc*AA10 and *Sc*CBM2) or 2.5 g/L (*Sc*LPMO10C). The experiment with both *Sc*AA10 and *Sc*CBM2 in panel C contained 1 g/L of each protein.

Due to low protein yields, DSC could not be used for the engineered LPMO variants, which were therefore assessed using a thermoshift differential scanning fluorimetry (DSF) assay to determine *T*_*m*(app)_ in experiments that included the wild-type full-length enzyme, *Sc*AA10 and *Sc*CBM2 (Fig. S7). The melting temperatures derived from DSC were slightly higher (+1–2 °C) than the values obtained by DSF, but, generally, the *T*_*m*(app)_ values followed the same trend (Table 2). While SR displayed a very similar *T*_*m*(app)_ compared to WT, the two other variants (SL and EL) showed a somewhat lower *T*_*m*(app)_. In all cases, removal of the copper by adding EDTA reduced the *T*_*m*(app)_ by ∼9 °C, except for *Sc*CBM2, which has no copper site, and *Sc*AA10 domain for which the difference in *T*_*m*(app)_ was as large as ∼15 °C. The latter is remarkable and underpins the notion that the stability of this domain is affected by interactions with the linker and/or CBM.

**Table 2.**
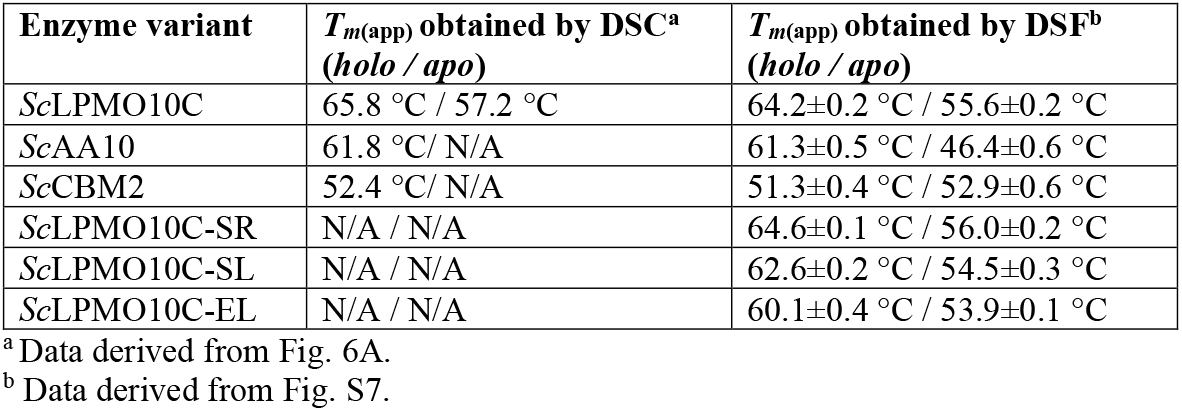
Apparent melting temperatures measured by DSC and DSF.

| **Enzyme variant** | **$T_{m(app)}$ obtained by DSC<sup>a</sup>**<br>**(*holo* / *apo*)** | **$T_{m(app)}$ obtained by DSF<sup>b</sup>**<br>**(*holo* / *apo*)** |
| --- | --- | --- |
| ScLPMO10C | 65.8 °C / 57.2 °C | 64.2±0.2 °C / 55.6±0.2 °C |
| ScAA10 | 61.8 °C / N/A | 61.3±0.5 °C / 46.4±0.6 °C |
| ScCBM2 | 52.4 °C / N/A | 51.3±0.4 °C / 52.9±0.6 °C |
| ScLPMO10C-SR | N/A / N/A | 64.6±0.1 °C / 56.0±0.2 °C |
| ScLPMO10C-SL | N/A / N/A | 62.6±0.2 °C / 54.5±0.3 °C |
| ScLPMO10C-EL | N/A / N/A | 60.1±0.4 °C / 53.9±0.1 °C |

<sup>a</sup> Data derived from Fig. 6A.
<sup>b</sup> Data derived from Fig. S7.

Encouraged by the apparent ability of *Sc*LPMO10C to withstand heat treatment, we then carried out a final test of how the linkers affect protein thermal stability. The various proteins were boiled for 0–120 min prior to starting LPMO reactions by adding Avicel and ascorbic acid (Fig. 7 and Fig. S8A) or carrying out a binding assay (for *Sc*CBM2; Fig. S8B). Of note, the residual activity assays are not fully quantitative since LPMO catalysis in these standard assays is limited by the *in situ* generation of the co-substrate, H_2_O_2_, and not linearly dependent on the concentration of active enzyme (Stepnov *et al*, 2022b; Filandr *et al*, 2020). Fig. 7 shows that for all linker variants, enzyme inactivation was not prominent after 60 minutes of pre-incubation and three of the four variants behave similarly (Fig. 7B). In line with its higher *T*_*m(app)*_ value, SR, containing a linker with a sequence that is very different from all others, stands out by showing barely any signs of inactivation after 60 minutes (Fig. 7B).

**Figure 7.**
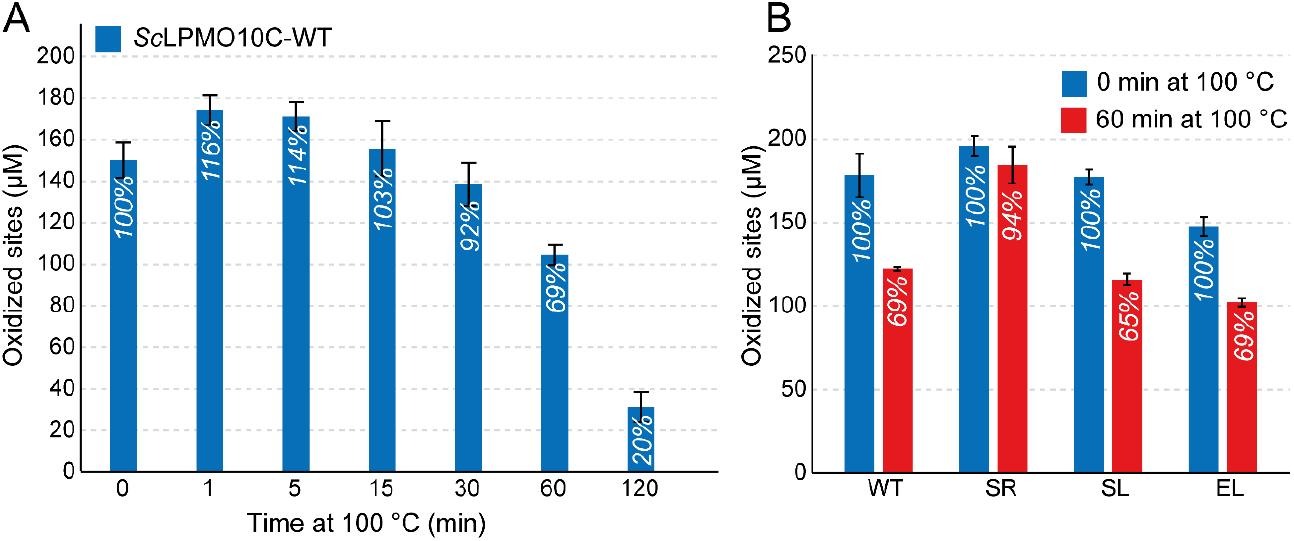
Residual activity after boiling. (A) Cellulose oxidation by *Sc*LPMO10C that had been pre-exposed to boiling for various amounts of time. (B) Enzyme activity of the four linker types in reactions that were not heat exposed, and in reactions where the enzymes were boiled for 60 min. The reactions were carried out in 50 mM sodium phosphate buffer (pH 6.0) in a thermomixer set to 30 °C and 800 rpm. The error bars show ±S.D. (n=3).

Binding studies with heat pretreated *Sc*CBM2 showed that the unfolding of this domain is reversible, since the ability to bind cellulose was retained (Fig. S8B). Studies with *Sc*AA10 showed that this domain is more sensitive to heat treatment compared to the full-length enzyme, with almost no activity retained after 60 minutes of incubation (Fig. S8A). This adds to the notion that both the linker and CBM2 increase the structural integrity and stability of *Sc*LPMO10C.

## Discussion

Using a combination of SAXS, NMR and MD simulations we determined global structural features of full-length *Sc*LPMO10C with its wild-type linker (Figs. 2–3, S1–3 and Table 1). The present results are a considerable improvement over the previous model of this enzyme (Courtade *et al*, 2018), as we obtained a more accurate determination of the overall size in terms of *R*_*g*_ and *R*_*h*_, and a vastly improved coverage of the dynamic features of the linker region in the ps–ns time scale, which were probed by ^15^N relaxation. The data show a linker that is flexible but, at the same time, favors an extended conformation.

After using SAXS data to optimize the Martini force field we were able to generate conformational ensembles of *Sc*LPMO10C with four different linkers using MD simulations. The force field modification consisted of a 10% increase of the protein-water interactions, which is in accordance with the 6–10% range found by Thomasen *et al* (Thomasen *et al*, 2022) when optimizing the agreement between SAXS data and simulations of 12 IDPs and three multidomain proteins using the Martini force field. The results of the MD simulations (Table 1) did not show a correlation between linker length and the calculated radius of gyration of the complete protein, nor was there a good correlation between the calculated radius of gyration and the hydrodynamic radius determined by NMR. Next to revealing limitations of either the molecular dynamics simulations or the NMR-based measurements of *R*_*h*_, these observations suggest that variations in linker structure impact the spacing between the two domains.

Sequence-dependent structural variation in the linker was assessed by simulations with a simpler single-bead model to analyze a large collection of natural linkers, which is an approach shown to be useful for assessing the conformation of flexible proteins (Tesei *et al*, 2021). This analysis clearly showed that the *R*_*g*_ of the 164 natural linker sequences analyzed depends mostly on the number of amino acids and not on their identity, although the model encodes sequence specificity. This observation finds explanation in the low complexity of the linker sequences (Fig. 4E), which are predominantly rich in glycine, threonine and proline. This finding contrasts recent work on the conformational buffering of disordered linkers in the adenovirus early gene 1A (E1A) protein (González-Foutel *et al*, 2022). The end-to-end distances of E1A linkers appear to be under functional selection and variations in sequence length and composition are compensated for to maintain an optimal end-to-end distance that maximizes binding to the E1A interaction partner (the retinoblastoma protein) (González-Foutel *et al*, 2022).

Importantly, this analysis of natural linkers leads to the conclusion that the above-mentioned lack of correlation likely is not due to variation in linker structure and, thus, may be due to variation in interactions that involve the two protein domains. Such interactions also became apparent from the functional characterization of the enzyme variants, as discussed below.

We assessed multiple potential functional roles of the linker. Measurements of catalytic performance (Fig. 5A, B) showed that the two variants containing equally long linkers with very different sequences have equal performance. The variants with a shorter or longer linker showed signs of increased enzyme inactivation, which, in the case of LPMOs, is indicative of less optimal enzyme-substrate interactions. It is conceivable that linker length affects the product distribution, since the linker restricts the area that the catalytic domain can reach when anchored to the substrate through the appended CBM. However, differences in product distributions were not detected (Fig. 5C, D). Assessment of thermal stability by DSF showed that the linker variant with wild-type like catalytic performance, SR, was also as equally stable as the wild-type, whereas the two poorer performing variants, EL and SL, demonstrated slightly reduced apparent *T*_m_ values. Importantly, all unfolding curves showed a single transition, which indicates that the AA10 and CBM2 domains interact in a manner that is mediated by the linker. The interactions between the domains and the role of the linker are supported by the DSC studies of the wild-type enzyme (Fig. 6), which show (1) that the full-length enzyme has a single unfolding transition with an apparent *T*_m_ that is higher than the *T*_m_ of the unfolding transitions observed for each of the individual domains and (2) that a mixture of the two domains has two folding transitions yielding an unfolding curve that represents the sum of the unfolding curves for the individual domains (Fig. 6C). Of note, the final experiments, in which we subjected the LPMO to boiling prior to an activity assay (Fig. 7 & S7), showed that the isolated catalytic domain, *Sc*AA10, is less stable than the full-length enzyme, adding to the notion that the presence of the linker and the CBM add to overall enzyme stability.

Our observations suggest that evolutionary pressure appears to shape linker length in LPMOs with two-domain architectures similar to *Sc*LPMO10C, rather than influencing amino acid composition. In line with our observations, a study of linkers in cellulases also indicated a consistent linker length (34–35 amino acids) in bacterial GH6 enzymes with CBM2s independent of the location of the CBM (N- or C-terminal) (Sammond *et al*, 2012).

Although our assessment of four linker variants indicates that linker length is most important and, in this study, a length of 34 residues was shown to be optimal for *Sc*LPMO10C, our analysis of 164 linkers shows that there is no universally optimal functional linker for LPMOs with the domain organization of *Sc*LPMO10C (i.e., AA10-linker-CBM2). The linker in *Sc*LPMO10C acts as a flexible spacer, giving the protein a dumbbell shape (Fig. 1 and 3A) where the individual domains are kept apart while preserving independent motions. At the same time, all other data show that linker-mediated domain interactions play a role. This is supported by a previous study on an LPMO produced by *Hypocrea jecorina* (*Hj*LPMO9A), which showed that specific portions of the linker interact with the catalytic domain (Hansson *et al*, 2017). An ideal functional linker would have a length and composition that favors stabilizing interactions either between the linker and domains or between the domains themselves. This is underscored by the observation of *Sc*LPMO10C resilience, maintaining activity even after boiling (Fig. 7). Moreover, the variant with the serine-rich (SR) linker with the same length as the wild-type exhibited even higher resistance to boiling, hinting at inherent stabilizing interactions for these linker lengths.

Linker variability may also relate to hitherto undetected variation in LPMO substrate specificity. Cellulosic substrates exist in multiple forms and cellulose crystals have multiple faces with differing morphologies (Zugenmaier, 2001). Our activity data (Fig 5A, B) show that the linker length affects how well the LPMO interacts with its substrate. It is thus conceivable that natural variation in LPMOs linker length reflects that these LPMOs have different optimal cellulose substrates.

This study found that, despite being extended and flexible, the linker affects interactions between the domains and with the substrate (as shown by the different stabilities under turnover conditions) and structural stability (as shown by the analysis of unfolding and thermal stability). So, while structural data seem to show that the linker is “just” an extended tether, the functional data show that the linker is more than that. These insights carry pivotal implications for biotechnological applications. Beyond traditional biomass degradation, where LPMOs already have recognized potential (Harris *et al*, 2014; Johansen, 2016), there is an emerging interest in the role of LPMOs in cellulose defibrillation processes for nanocellulose production. In this context, the presence of a CBM and linker has been show to affect the carboxyl content in cellulose nanofibrils produced using LPMOs (Koskela *et al*, 2019). Future endeavors in linker-oriented protein engineering could leverage these insights to develop or optimize enzymes tailored for specific applications.

## Materials and methods

### Cloning, expression, and purification of ScLPMO10C and linker variants

Wild-type *Sc*LPMO10C and its truncated versions (*Sc*AA10 and *Sc*CBM2) were cloned and produced as previously described (Forsberg *et al*, 2014; Courtade *et al*, 2018) and loaded with copper according to the protocol described by (Loose *et al*, 2014). The pRSET B expression vector harboring *Sc*LPMO10C (UniProtID: Q9RJY2) (Forsberg *et al*, 2014) was used as a starting point to replace the linker region (residue 229-262) with i) a 34 amino acid poly-serine linker from *Cellvibrio japonicus* LPMO10A (*Cj*LPMO10A; UniProtID: B3PJ79, residues 217-250) (Forsberg *et al*, 2016), ii) a 20 amino acid linker from *Streptomyces scabies* LPMO10B (*Ssc*LPMO10B; UniProtID: C9YVY4, residue 238-257) (Stepnov *et al*, 2022a), and iii) an extended linker of 59 amino acids from *Caldibacillus cellulovorans* LPMO10A (*Cc*LPMO10A; UniProtID: Q9RFX5 residues 226-284). The sequences of these linkers are provided in Figure 1 and Table S1. DNA encoding for the alternative linkers was derived from gene sequences that were codon optimized for expression in *Escherichia coli*. The pRSET B_*sclpmo10c-wt* vector was amplified using a forward primer annealing to the start of the *cbm2* sequence (F-primer; 5’ GGTTCGTGTATGGCCGTCTATA 3’) and a reverse primer which anneals to the end of the *aa10* sequence (R-primer; 5’ GTCGAAAACCACATCGGAGCAA 3’), thereby amplifying the entire 2.8 kb vector excluding the 102 nucleotides encoding the linker. The amplified vector was gel purified (1% agarose) and used for In-Fusion® HD cloning (Clontech, Mountain View, CA, USA) with amplified and gel purified (2% agarose) linker inserts. The size of the inserts varied between 90 and 207 nucleotides. After fusing linker inserts to the linearized vector, One Shot™ TOP10 chemically competent *E. coli* cells (Invitrogen) were transformed. Positive transformants were subsequently verified by Sanger sequencing (Eurofins GATC, Cologne, Germany).

Expression of ^13^C- and ^15^N-labeled *Sc*LPMO10C for chemical shift assignment and ^15^N relaxation measurements was performed using the XylS/*Pm* expression system (Courtade *et al*, 2017b). The pJB_SP_*sclpmo10c-wt* vector was constructed as previously described (Courtade *et al*, 2017b), harboring *Sc*LPMO10C downstream of a pelB signal peptide.

For regular protein production, T7 Express competent *E. coli* (New England Biolabs catalog number 2566) were transformed using a heat-shock protocol and grown at 37 °C on LB-agar plates. All media were supplemented with 100 μg/ml ampicillin. Pre-cultures were made in shaking flasks by inoculating 5 mL LB medium (10g/L tryptone, 5g/L yeast extract, 5g/L NaCl) with recombinant cells, followed by incubation at 30 °C and 225 rpm overnight. Main cultures were made by inoculating 500 mL 2×LB medium (20 g/L tryptone, 10 g/L yeast extract, 5 g/L NaCl) with 1% pre-culture, followed by incubation in a LEX-24 bioreactor (Epiphyte3, Toronto, Canada) using compressed air (20 psi) for aeration and mixing. The SL, SR and EL variants were incubated at 30 °C for 24 hours, whereas the wild-type variant was incubated at 30 °C to OD_600nm_ ≈ 0.8. The latter culture was cooled on ice for 5 min, induced with 0.1 mM *m*-toluic acid, and further incubated at 16 °C for 20 h.

For the production of isotopically labeled protein, main cultures were made by inoculating 500 mL M9 medium (6 g/L Na_2_HPO_4_, 3 g/L KH_2_PO_4_, 0.5 g/L NaCl) supplemented with (^15^NH_4_)_2_SO_4_ (98% ^15^N), 4 g/L unlabeled or ^13^C-labeled glucose (98% uniformly labeled with ^13^C), 10 mL Bioexpress Cell Growth Media (Cambridge Isotope Laboratories, Tewksbury, MA, USA), 5 mL Gibco™ MEM Vitamin Solution (100x), and trace metal solution (300 mg/mL MgSO_4_, 2 mg/L ZnSO_4_, 10 mg/L FeSO_4_, 2 mg/L CuSO_4_, and 20 mg/L CaCl_2_) with 1% pre-culture, followed by incubation at 30 °C to OD_600nm_ ≈ 0.8. The culture was cooled on ice for 5 min, induced with 0.1 mM *m*-toluic acid, and further incubated at 16 °C for 20 h.

Cells were harvested by centrifugation for 5 min at 5500×g, and 4 °C, and periplasmic fractions were prepared by the osmotic shock method, as follows: the pellet was resuspended in 30 mL spheroplast buffer (pH 7.5, 100 mM Tris-HCL, 500 mM sucrose, 0.5 mM EDTA) with a tablet of cOmplete™ ULTRA protease inhibitor (Roche), followed by centrifugation for 10 minutes at 6150×g and 4°C. The pellet was incubated at room temperature for 10 minutes, prior to resuspension in 25 mL ice-cold water with another tablet of cOmplete™ ULTRA protease inhibitor (Roche), followed by centrifugation for 30 minutes at 15000×g and 4 °C. The supernatant was filtered through a 0.2 μm pore size filter prior to protein purification. The proteins were purified by loading periplasmic extracts in Buffer A (50 mM Tris-HCl, pH 8.5) onto a 5 mL HiTrap DEAE Sepharose FF anion exchanger (Cytiva, Marlborough, MA, USA) connected to an ÄKTA™ Pure FPLC system (Cytiva). LPMOs were eluted by applying a linear salt gradient towards 50 mM Tris-HCl, pH 8.5, 1 M NaCl (Buffer B), at a flow rate of 5 mL/min. The LPMOs started to elute at 10-12% Buffer B. The fractions containing LPMO were pooled and concentrated using ultrafiltration spin-tubes (10 kDa cut-off, Sartorius). The protein-containing fractions were analyzed by SDS-PAGE.

Before SAXS measurements, proteins were further purified by size-exclusion chromatography. Samples were loaded onto a HiLoad® 16/600 Superdex® 75 pg size-exclusion column (Cytiva), using 50 mM Tris-HCl, pH 7.5, 200 mM NaCl as running buffer and a flowrate of 1 mL/min.

### SAXS

*Sc*LPMO10C was measured by SAXS at the P12 beamline at DESY (Blanchet *et al*, 2015). The scattering intensity, I(q), was measured as a function of q = 4π sin θ/λ, where q is the scattering vector, 2θ is the scattering angle, and λ is the X-ray wavelength (1.24 Å at P12). Buffer backgrounds were measured before and after each sample and all measurements were performed at 8 °C. The initial data reduction including azimuthal averaging, frame averaging, and background subtraction was performed using the PRIMUS program (Konarev *et al*, 2003) that is part of the ATSAS software (Franke *et al*, 2017). Data was converted into the usual units of scattering cross section per unit volume, cm^*−*1^, using the well-known scattering cross sections of, respectively, H_2_O and bovine serum albumin as external standards (Orthaber *et al*, 2000; Mylonas & Svergun, 2007). Further, the data was logarithmically rebinned with a bin factor of 1.02 using the WillItRebin program (Pedersen *et al*, 2013). Pair-wise distance distribution functions were calculated using the Bayesian Indirect Fourier Transformation (BIFT) algorithm implemented in BayesApp (Hansen, 2012).

### NMR

All NMR spectra were recorded in an NMR buffer (25 mM sodium phosphate and 10 mM NaCl, pH 5.5) containing 10% D_2_O (or 99.9% D_2_O for ^13^C-detected experiments; see below) at 25 °C using a Bruker Ascend 800 MHz spectrometer with an Avance III HD (Bruker Biospin) console equipped with a 5 mm z-gradient CP-TCI (H/C/N) cryogenic probe, at the NV-NMR-Centre/Norwegian NMR Platform at the Norwegian University of Science and Technology (NTNU). NMR data were processed and analyzed using Bruker TopSpin version 3.5, and Protein Dynamic Center software version 2.7.4 from Bruker BioSpin.

We performed a new chemical shift assignment of full-length *Sc*LPMO10C using previously published (Courtade *et al*, 2017a) backbone assignments for CBM2 and AA10 as the starting point, together with newly acquired data from the following experiments. Resonances for non-proline amino acids, i.e., possessing amide protons, were assigned by using ^15^N-HSQC, HNCA, HNCO and CBCA(CO)NH. The ^13^C-detected experiments CON, CANCO and CACO were used to assign backbone resonances for prolines in the linker region (Bermel *et al*, 2005).

Secondary structure elements in the linker region were analyzed using the web-based version of the TALOS-N software (http://spin.niddk.nih.gov/bax/software/TALOS-N/) (Shen & Bax, 2013) using the ^13^C and ^15^N chemical shifts.

Nuclear spin relaxation rates *R*_1_ and *R*_2_, and heteronuclear ^1^H-^15^N NOE measurements of amide ^15^N were recorded as pseudo-3D spectra where two frequency dimensions correspond to the amide ^1^H and ^15^N chemical shifts, respectively, and the third dimension is made up of variable relaxation time delays. For *R*_1_, the time points were 100, 200, 500, 1000, 1500, 2000, 3000, and 4000 ms. For *R*_2_, the time points were 17, 34, 102, 170, 204, and 238 ms. For ^1^H-^15^N NOE, two 2D planes were recorded, one with and one without pre-saturation. The generalized order parameter, *S*^2^, was obtained using reduced spectral density mapping (Fushman, 2002).

To calculate the translational diffusion coefficients, *D*_*t*_, of WT, SR, SL and EL we used a pulsed-field gradient stimulated-echo sequence with 3-9-19 water suppression (stebpgp1s19) (Price *et al*, 2002). A linear gradient ramp of 32 points in the range 2-95% of the gradient strength (where 100% corresponds to 53.5 G/cm) was used, and the gradient length and delay between gradients were set to Δ = 2 ms and δ = 150 ms, respectively. While it is in principle possible to calculate the absolute hydrodynamic radius, *R*_*h*_, by using the Stokes-Einstein equation (Edward, 1970), this requires accurate knowledge of the viscosity of the sample solution, which is difficult to measure. An alternative approach for calculating *R*_*h*_ consists of using a reference molecule in the solution, with a known *R*_*h*_. Here, 1% dioxane was used as a reference because it does not interact with proteins, and its *R*_*h*_ is known (*R*_*h,ref*_ = 2.12 Å) (Wilkins *et al*, 1999). The diffusion coefficient (*D*_*t,ref*_) of dioxane was thus used to determine the *R*_*h*_ of the proteins:

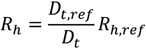

### Coarse-grained simulations using the Martini force field

The coarse-grained Martini version 3.0.beta.4.17 force field was used in combination with GROMACS 2018.2 (Abraham *et al*, 2015) to simulate *Sc*LPMO10C with different linkers. The constructs were coarse-grained using the Martinize2 program (de Jong *et al*, 2013). An elastic network model (Periole *et al*, 2009) was used to constrain the overall structure of the AA10 and CBM2 domains, but it was excluded from the linker region. The simulation box was filled with water beads and the system was neutralized with beads corresponding to Na^+^ and Cl^-^ ions to an ionic strength of 0.15 M. The complex was energy-minimized using a steepest-descent algorithm (100 steps, 0.03 nm max step size) prior to being relaxed for 1 ns, with a time step of 5 fs, using the velocity-rescale (v-rescale) thermostat (Bussi *et al*, 2007), Parrinello-Rahman barostat (Parrinello & Rahman, 1981) and Verlet cutoff scheme (Páll & Hess, 2013). Simulations were performed on the relaxed models with a time step of 20 fs, using the v-rescale thermostat, Parrinello-Rahman barostat and Verlet cutoff scheme in the isothermal-isobaric (NPT) ensemble.

Proteins simulated using Martini force fields have been observed to aggregate, possible due to protein-protein interaction strengths being too high relative to protein-water interactions (Larsen *et al*, 2020; Thomasen *et al*, 2022),, leading to mismatches between experimental data and simulations. To circumvent this problem we followed the approach described by Larsen et al (Larsen *et al*, 2020) to modify the Martini version 3.0.beta.4.17 forcefield to improve the fit between SAXS data and small-angle coarse-grained simulations. In summary, the interaction strength (i.e., ε parameter in the Lennard–Jones potential) between protein and water beads in the Martini force field was multiplied by a factor *λ*, ranging from 1.0 (unchanged) to 1.20 (20% increase of the protein-water interaction strength). Simulations for each value of *λ* were run for 10 μs and frames were written every 1 ns for each trajectory. To facilitate calculation of SAXS profiles, each of the 10,000 frames per CG simulations was used to reconstruct atomistic models by using the Backward program (Wassenaar *et al*, 2014).

### Calculating SAXS profiles from conformational ensembles

The implicit solvent SAXS calculation program Pepsi-SAXS version 3.0 (Grudinin *et al*, 2017) was used to calculate SAXS profiles from the conformational ensemble (i.e., the atomic coordinates of the 10,000 frames) of *Sc*LPMO10C. Input for Pepsi-SAXS comprises atomic coordinates, experimental SAXS profiles, and four parameters that can either be provided directly or determined by the program while fitting between the calculated and experimental SAXS profiles: the scale of the profiles, I(0), a constant background, *cst*, the effective atomic radius, *r*_0_, and the contrast of the hydration layer, δ_ρ_. The approach used here was based on the methods previously described by Larsen et al (Larsen *et al*, 2020) and Pesce et al (Pesce & Lindorff-Larsen, 2021). The parameter values were set to *I*(0) = 1.0, *cst* = 0.0, *r*_0_ = 1.65 Å and *δ*_*ρ*_ = 3.34 *e*/*nm*^3^. Then, the iBME (iterative Bayesian Maximum Entropy) method (Pesce & Lindorff-Larsen, 2021) was used to iteratively rescale and shift the calculated SAXS profiles, while reweighting the conformational ensemble to fit a global value of *I*(0) and *cst*.

The implicit solvent SAXS calculation program CRYSOL (Svergun *et al*, 1995) was used to calculate a SAXS profile for the structure (PDB: 4OY7) of *Sc*AA10. The search limits for the fitting parameters *dro* (optimal hydration shell contrast), *Ra* (optimal atomic group radius) and *ExVol* (relative background) were set to [0.000 – 0.075], [1.40 – 1.80] and [23744 – 27594], respectively. The parameters for maximum order of harmonics were set to 15, the order of Fibonacci grid to 17, and the electron density of the solvent was set to 0.33 e/Å^3^.

### Linker simulations using a single-bead model

To find linker sequences similar to the linker in wild-type *Sc*LPMO10C, we used *phmmer* in the HMMER webserver (Potter *et al*, 2018) to search the UniProt KB Reference Proteomes (Bateman *et al*, 2015) database against the Pfam (Finn *et al*, 2014) profile of *Sc*LPMO10C, i.e. AA10-linker-CBM2. Using this approach, we found 160 unique sequences with the same domain architecture as *Sc*LPMO10C. For each retrieved sequence, the AA10 and CBM2 domains were assigned using *hmmscan* and the linker region was assigned as the amino acids in between the domains.

We added the four linker sequences of the wild-type enzyme, SR, SL and EL to the retrieved sequences, and simulated the 164 sequences at 298 K, I=0.1 M and pH=6.5 using a model where each amino acid in the sequence is represented by a single bead placed at the Cα coordinates. The model has previously been optimized and validated against experimental SAXS and paramagnetic relaxation enhancement (PRE) NMR data of 48 IDPs of sequence length ranging from 24 to 334 residues (Tesei *et al*, 2021). In particular, for the amino acid specific “stickiness” parameters, we used the M1 set proposed by Tesei et al. (Tesei *et al*, 2021). With the aim of mimicking end-capped proteins, we didn’t modify the charges of the terminal residues, as opposed to other implementations of the model where the charge of the terminal carboxylate and amino groups are considered. Langevin simulations were performed using HOOMD-blue v2.9.3 (Anderson *et al*, 2020) using a time step of 5 fs, a friction coefficient of 0.01 ps^-1^, and a cutoff of 4 nm for both Debye-Hückel and nonionic nonbonded interactions. Each simulation started from the fully extended chain placed in a cubic box of 200 nm and we extensively sampled chain conformations for a total simulation time of 14 μs.

The persistence length was calculated by fitting the autocorrelation function of the bond vectors along the sequence to a single exponential decay:

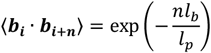

where ***b***_***i***_ and ***b***_***i+n***_ are bond vectors separated by *n* bonds, *l*_*b*_ = 0.38 nm (the equilibrium bond length), and *l*_*p*_ is the persistence length.

The scaling exponent was calculated by fitting the long-distance region, |*i − j*| > 10, of the average intramolecular pairwise distances, *R*_*ij*_, to the following equation (Shrestha *et al*, 2021)

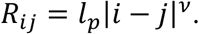

The R_g_ values of the linkers were estimated from the sequence length via the following equation (Zheng *et al*, 2020)

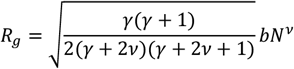

where *γ* = 1.1615, *b* = 0.55 nm, and ν is the average scaling exponent for the 164 linker sequences.

### LPMO activity assays

Enzyme activity assays were set up using 10 g/L Avicel, 0.5 μM LPMO, 1 mM ascorbic acid or 1 mM gallic acid (in 100% DMSO) in 50 mM sodium phosphate buffer, pH 6.0, in the absence or presence of 100 μM H_2_O_2_, and were incubated for up to 48 h in a thermomixer set to 30 °C. Aliquots were taken at various time points and vacuum filtered through a 0.45 μm membrane to separate the solid fraction. The supernatants (i.e. the soluble fractions) were further treated with *Tf*Cel6A (E2; (Spezio *et al*, 1993)) to convert longer oxidized cello-oligosaccharides to oxidized dimers and trimers, which can be quantified by HPAEC-PAD, using in-house made standards (Bissaro *et al*, 2016). To determine the total concentration of oxidized products (i.e., soluble and insoluble), the LPMOs were inactivated by boiling the reaction mixture for 15 min at 100 °C. Subsequently, the reaction mixtures were cooled on ice and diluted in buffer to a final concentration of 2 g/L Avicel including 5 μM *Tf*Cel6A and 2 μM *Tf*Cel5A (Irwin *et al*, 1993), followed by 48-h incubation at 50 °C in an Eppendorf Thermomixer with shaking at 800 rpm. As a result of this procedure, all oxidized sites are solubilized as oxidized dimers and trimers, which were quantified by HPAEC-PAD (see below).

Product distribution profiles were assessed by HPAEC-PAD using samples of LPMO-generated soluble products that had not been treated with cellulases. The peak areas (nC×min) of oxidized LPMO products were used to calculate apparent ratios of oxidized oligosaccharides with different degrees of polymerization (DP2-DP6).

### Residual activity and cellulose binding of heat-treated proteins

To test whether heat-treated and potentially refolded LPMOs retained their functional properties, solutions of WT and *Sc*AA10 at a concentration of 2 μM in 50 mM sodium phosphate buffer, pH 6.0, were boiled for 0-120 min and then rapidly cooled on ice prior to starting the LPMO reaction by addition of reductant (1 mM ascorbic acid) and substrate (10 g/L Avicel). After incubation of the reactions for 24 h at 30 °C, *Tf*Cel6A was added to filtrated supernatants and the oxidized dimers and trimers were quantified using HPAEC-PAD to determine residual LPMO activity.

The A_280_ of a solution with 0.2 g/L CBM2 in 50 mM sodium phosphate buffer, pH 6.0, was measured prior to boiling this solution for 0-120 min. After cooling the samples on ice, the proteins were filtrated using a 96-well filter plate (Millipore, Burlington, MA, USA) to remove precipitated protein and the A_280_ was remeasured before starting the binding assay. Importantly, the loss in protein concentration due to precipitation was barely noticeable. For the binding assay, reactions were set up containing 0.1 g/L of heat-pretreated protein and 10 g/L Avicel in 50 mM sodium phosphate buffer, pH 6.0, and incubated at 800 rpm and 22 °C. After 1 h of incubation the samples were filtrated and A_280_ was measured again to estimate the concentration of unbound CBM2.

Control reactions containing (i) 2 mM ascorbic acid in buffer, or (ii) 2 μM *Sc*LPMO10C and 2 mM ascorbic acid in buffer were boiled for 15 min prior to starting reactions by adding *Sc*LPMO10C and Avicel (i) or Avicel only (ii). In the end, all reactions contained 10 g/L Avicel, 1 mM ascorbic acid, 1 μM *Sc*LPMO10C in 50 mM sodium phosphate buffer, pH 6.0. The reactions were incubated overnight followed by filtration, treatment with *Tf*Cel6A and quantification of oxidized products by HPAEC-PAD.

### High-performance anion-exchange chromatography with pulsed amperometric detection (HPAEC-PAD)

HPAEC-PAD analysis of LPMO-generated products was carried out using an ICS-5000 system from Dionex (Sunnyvale, CA) equipped with a disposable electrochemical gold electrode as previously described (Westereng *et al*, 2013). Samples of 5 μL were injected onto a CarboPac PA200 (3×250 mm) column operated with 0.1 M NaOH (eluent A) at a flow rate of 0.5 mL/min and a column temperature of 30 °C. Elution was achieved using a stepwise gradient with increasing amounts of eluent B (0.1 M NaOH + 1 M NaOAc), as follows: 0–5.5% B over 3 min; 5.5–15% B over 6 min; 15–100% B over 11 min; 100–0% B over 0.1 min; and 0% B (reconditioning) for 5.9 min. Chromatograms were recorded and analyzed by peak integration using Chromeleon 7.0 software.

### Apparent melting temperatures

A Nano-Differential Scanning Calorimeter III (Calorimetry Sciences Corporation, Lindon, USA) was used to determine *T*_*m*(*app*)_ of *Sc*LPMO10C and its individual domains. Solutions containing 1 mg/mL (*Sc*AA10 and *Sc*CBM2) or 2.5 mg/mL (full-length) protein in 50 mM sodium phosphate buffer, pH 6.0, (filtered and degassed) were heated from 25 °C to 90 °C at 1 °C/min followed by cooling from 90 °C to 25 °C at the same rate. At least three cycles (i.e., three heating and three cooling steps) were recorded for each run. Buffer baselines were recorded and subtracted from the protein scans unless stated otherwise. The melting curve for the *apo*-form of *Sc*LPMO10C was obtained in the same manner by introducing 5 mM EDTA to both the protein sample and the control sample (i.e., sample lacking the enzyme). The data were analyzed with NanoAnalyze software (https://www.tainstruments.com).

For the engineered enzyme variants that did not efficiently express, a differential scanning fluorimetry assay based on the use of SYPRO orange (Thermoshift assay kit, Thermo Fisher Scientific) was used to minimize protein consumption (Huynh & Partch, 2015). The quantum yield of the SYPRO orange dye is significantly increased upon binding to hydrophobic regions of the protein that become accessible as the protein unfolds. The fluorescence emission was monitored using a StepOnePlus real-time PCR machine (Thermo Fisher Scientific). The apparent *T*_m_ was calculated as the temperature corresponding to the minimum value of the derivative plot (*−*d[relative fluorescence unit]/dT versus T; Fig. S7). Solutions containing 0.1 g/L protein and SYPRO orange (1×) in 50 mM sodium phosphate buffer, pH 6.0, were heated in a 96-well plate from 25 to 95 °C, over 50 min. For each protein, the experiment was carried out in quadruplicates (i.e., n = 4).

## Data availability

NMR chemical shift assignments have been deposited in the BioMagnetic Resonance Databank (BMRB) under the ID 27078. All the data, input files and code required to reproduce the simulation results reported in this article, including the protein ensembles, are available online at https://github.com/gcourtade/papers/tree/master/2023/ScLPMO10C-linkers.

## Acknowledgements

This work was funded by The Novo Nordisk Foundation, project numbers NNF18OC0055736 (to Z. F.) and NNF18OC0032242 (to G. C.), and by the Research Council of Norway through projects 226244, 269408 (to V.E.) and 262853 (to V.E.). Y. W., G. T. and K. L.-L. were supported by the BRAINSTRUC structural biology initiative from the Lundbeck Foundation. Part of the computations were performed on resources provided by Sigma2 - the National Infrastructure for High Performance Computing and Data Storage in Norway.

