## Supplementary Information for "The effect of linker conformation on performance and stability of a two-domain lytic polysaccharide monooxygenase"

**Table S1. Primary structure of the four linker variants of *Sc*LPMO10C.** Red and blue labelled sequences represent the AA10 and the CBM2 domains, respectively, and are the same in all variants. The black highlighted sequences show the four different linkers. Note that the first nine residues in the short linker (SL) are identical to the first part of the wild-type linker; for clarity the rest of this linker is shown in red face (on black background).

| **Enzyme name** | **$M_w$ (Da)** | **Protein sequence** |
|---|---|---|
| *Sc*AA10<br>(isolated AA10 catalytic domain) | 20,869 | HGVAMMPGSRTYLCQLDAKTGTGALDPTNPACQAALDQSGATALYNW<br>FAVLDSNAGGRGAGYVPDGTLCSAGDRSPYDFSAYNAARSDWPRTHL<br>TSGATIPVEYSNWAAHPGDFRVYLTKPGWSPTSELGWDDLELIQTVT<br>NPPQQGSPGTDGGHYYWDLALPSGRSGDALIFMQWVRSDSQENFFSC<br>SDVVFD |
| *Sc*LPMO10C-**WT**<br>(**W**ild-**T**ype linker) | 34,568 | HGVAMMPGSRTYLCQLDAKTGTGALDPTNPACQAALDQSGATALYNW<br>FAVLDSNAGGRGAGYVPDGTLCSAGDRSPYDFSAYNAARSDWPRTHL<br>TSGATIPVEYSNWAAHPGDFRVYLTKPGWSPTSELGWDDLELIQTVT<br>NPPQQGSPGTDGGHYYWDLALPSGRSGDALIFMQWVRSDSQENFFSC<br>SDVVFDGGNGEVTGIRGSGSTPDPDPTPTPTDPTTPPTHTGSCMAVY<br>SVENSWSGGFQGSVEVMNHGTEPLNGWAVQWQPGGGTTLGGVWNGSL<br>TSGSDGTVTVRNVDHNRVVPPDGSVTFGFTATSTGNDFPVDSIGCVA<br>P |
| *Sc*LPMO10C-**SR**<br>(**S**erine-**R**ich linker) | 34,287 | HGVAMMPGSRTYLCQLDAKTGTGALDPTNPACQAALDQSGATALYNW<br>FAVLDSNAGGRGAGYVPDGTLCSAGDRSPYDFSAYNAARSDWPRTHL<br>TSGATIPVEYSNWAAHPGDFRVYLTKPGWSPTSELGWDDLELIQTVT<br>NPPQQGSPGTDGGHYYWDLALPSGRSGDALIFMQWVRSDSQENFFSC<br>SDVVFDNGTGTGSSSSVASSVVSSVTSSSVASSVASSLSNGSCMAVY<br>SVENSWSGGFQGSVEVMNHGTEPLNGWAVQWQPGGGTTLGGVWNGSL<br>TSGSDGTVTVRNVDHNRVVPPDGSVTFGFTATSTGNDFPVDSIGCVA<br>P |
| *Sc*LPMO10C-**SL**<br>(**S**hortened **L**inker) | 33,186 | HGVAMMPGSRTYLCQLDAKTGTGALDPTNPACQAALDQSGATALYNW<br>FAVLDSNAGGRGAGYVPDGTLCSAGDRSPYDFSAYNAARSDWPRTHL<br>TSGATIPVEYSNWAAHPGDFRVYLTKPGWSPTSELGWDDLELIQTVT<br>NPPQQGSPGTDGGHYYWDLALPSGRSGDALIFMQWVRSDSQENFFSC<br>SDVVFDGGNGEVTGIKQPGNPSEPVPGSCMAVYSVENSWSGGFQGSV<br>EVMNHGTEPLNGWAVQWQPGGGTTLGGVWNGSLTSGSDGTVTVRNVD<br>HNRVVPPDGSVTFGFTATSTGNDFPVDSIGCVAP |
| *Sc*LPMO10C-**EL**<br>(**E**xtended **L**inker) | 37,126 | HGVAMMPGSRTYLCQLDAKTGTGALDPTNPACQAALDQSGATALYNW<br>FAVLDSNAGGRGAGYVPDGTLCSAGDRSPYDFSAYNAARSDWPRTHL<br>TSGATIPVEYSNWAAHPGDFRVYLTKPGWSPTSELGWDDLELIQTVT<br>NPPQQGSPGTDGGHYYWDLALPSGRSGDALIFMQWVRSDSQENFFSC<br>SDVVFDSGPIAYEFGDPREGGTMITPPPSGTTPTPTPTPTPTSTPTP<br>TPTPSVTPTVTPTSTPTPGSCMAVYSVENSWSGGFQGSVEVMNHGTE<br>PLNGWAVQWQPGGGTTLGGVWNGSLTSGSDGTVTVRNVDHNRVVPPD<br>GSVTFGFTATSTGNDFPVDSIGCVAP |

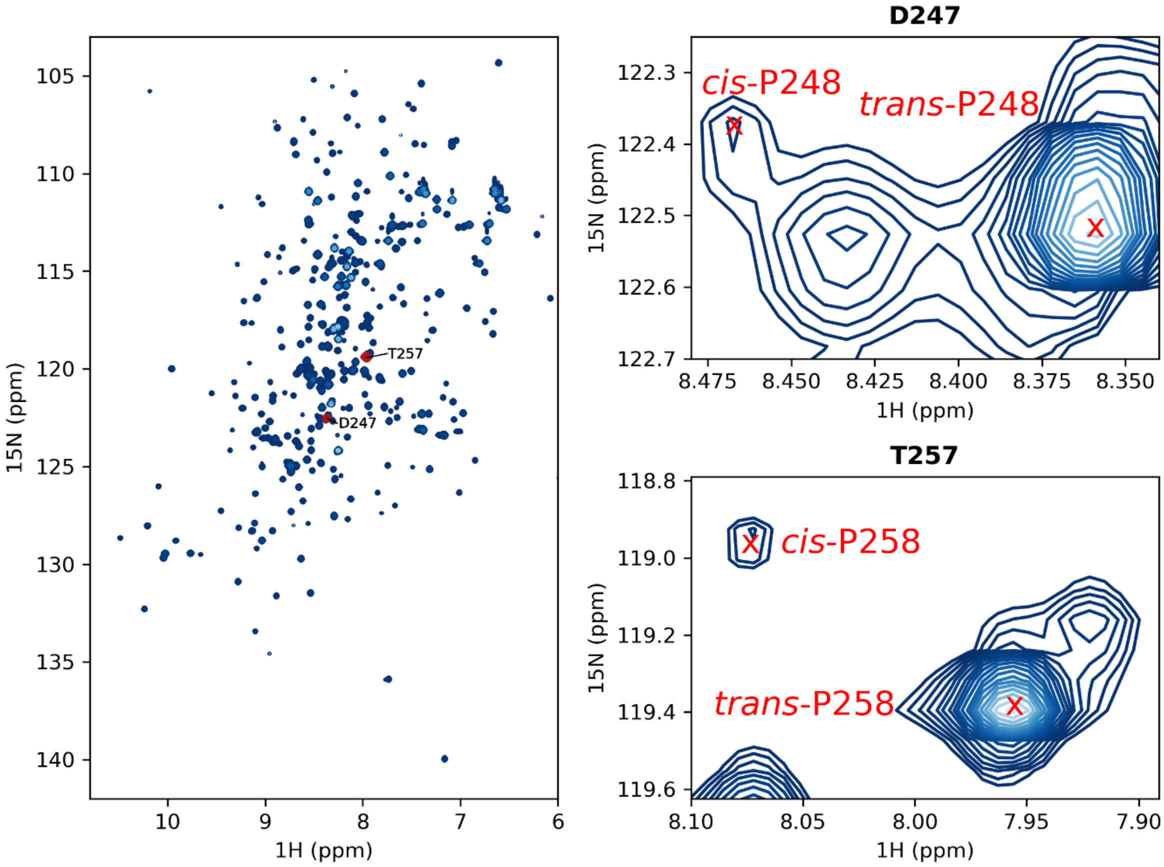


**Figure S1**. **Evidence of of *cis*-Pro bonds in *Sc*LPMO10C.** Left panel: 2D $^{1}$H-$^{15}$N HSQC spectrum of 80 μM $^{13}$C- and $^{15}$N-labeled *Sc*LPMO10C at pH 5.5 and 298 K. Resonances affected by *cis-trans* Pro isomerism of neighboring Pro residues (D247 and T257) are colored red. Right panels: Zoomed-in regions of the 2D $^{1}$H-$^{15}$N HSQC spectrum. The peaks corresponding to the major *trans*-Pro and minor *cis*-Pro populations are indicated. The *cis*-fraction was estimated by the peak intensities, *I*, as $I_{cis}/(I_{cis} + I_{trans})$.

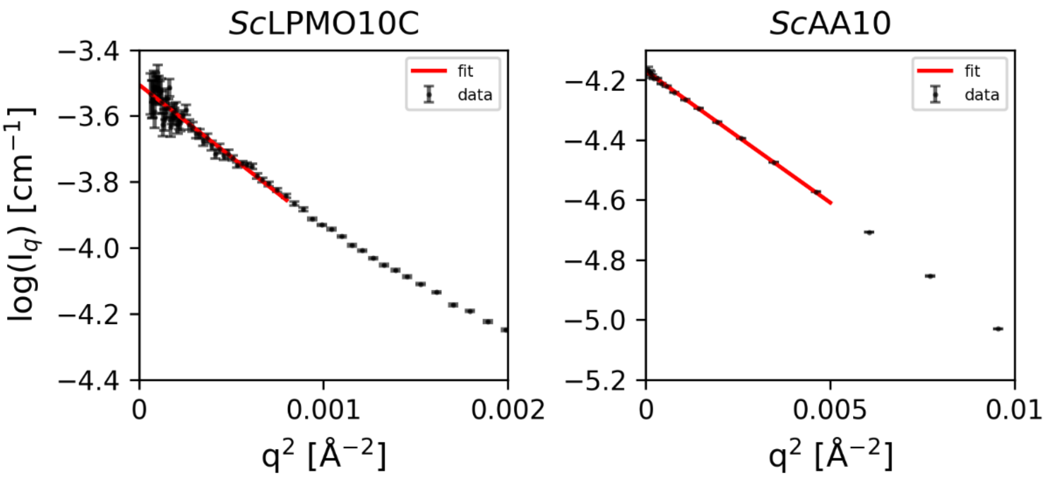


**Figure S2. Guinier plots.** Used to calculate the radii of gyration ($R_g$) from SAXS data for *Sc*LPMO10C and its catalytic domain (*Sc*AA10).

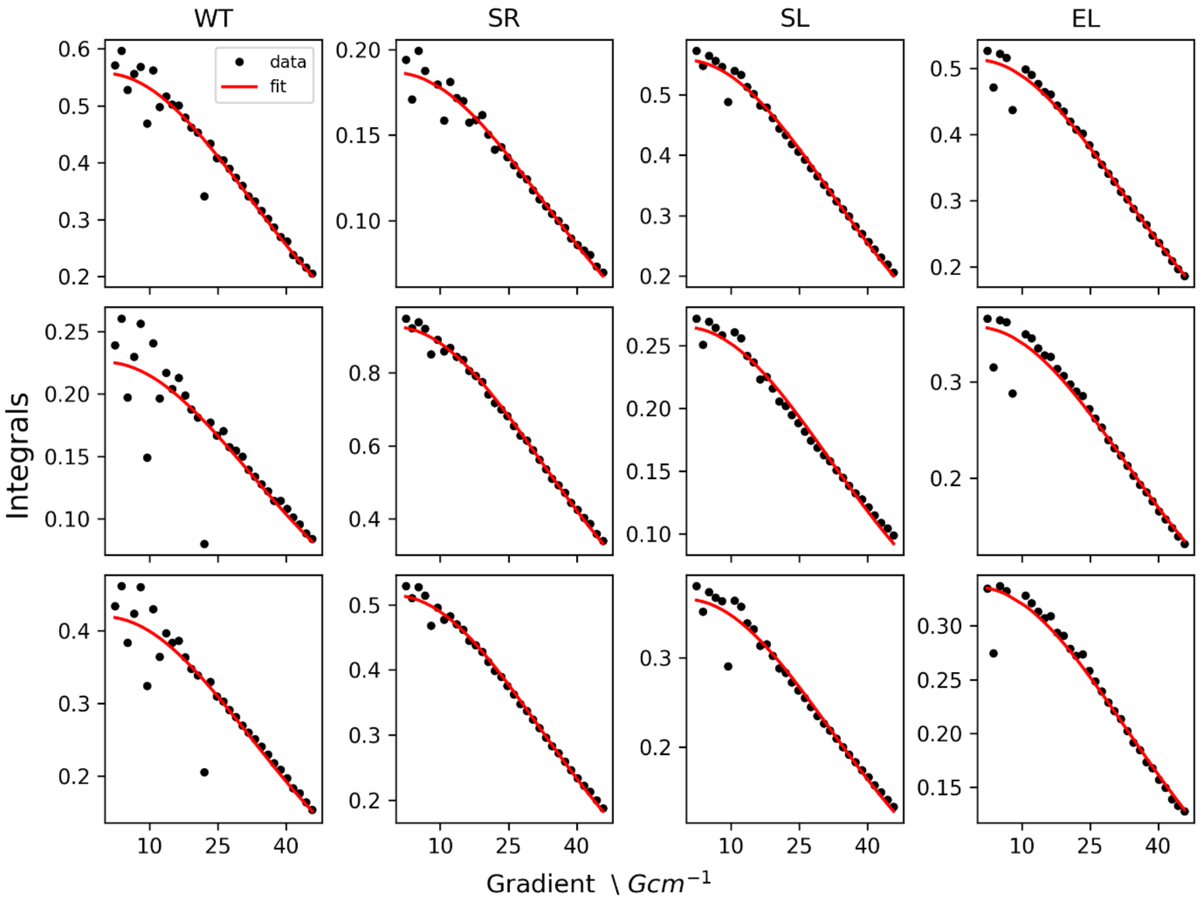


**Figure S3. Diffusion decay curves.** The graphs show integrals of three selected signals in the DOSY spectra of *Sc*LPMO10C and three linker variants, as a function of gradient strength. The fitted diffusion model is shown in red, and the estimated diffusion constants in $10^{-1}$ $m^2/s$ are, from top to bottom, WT: 1.14, 1.14, 1.13; SR: 1.13, 1.15, 16; SL: 1.15, 1.18, 1.17; EL: 1.12, 1.09, 1.07.

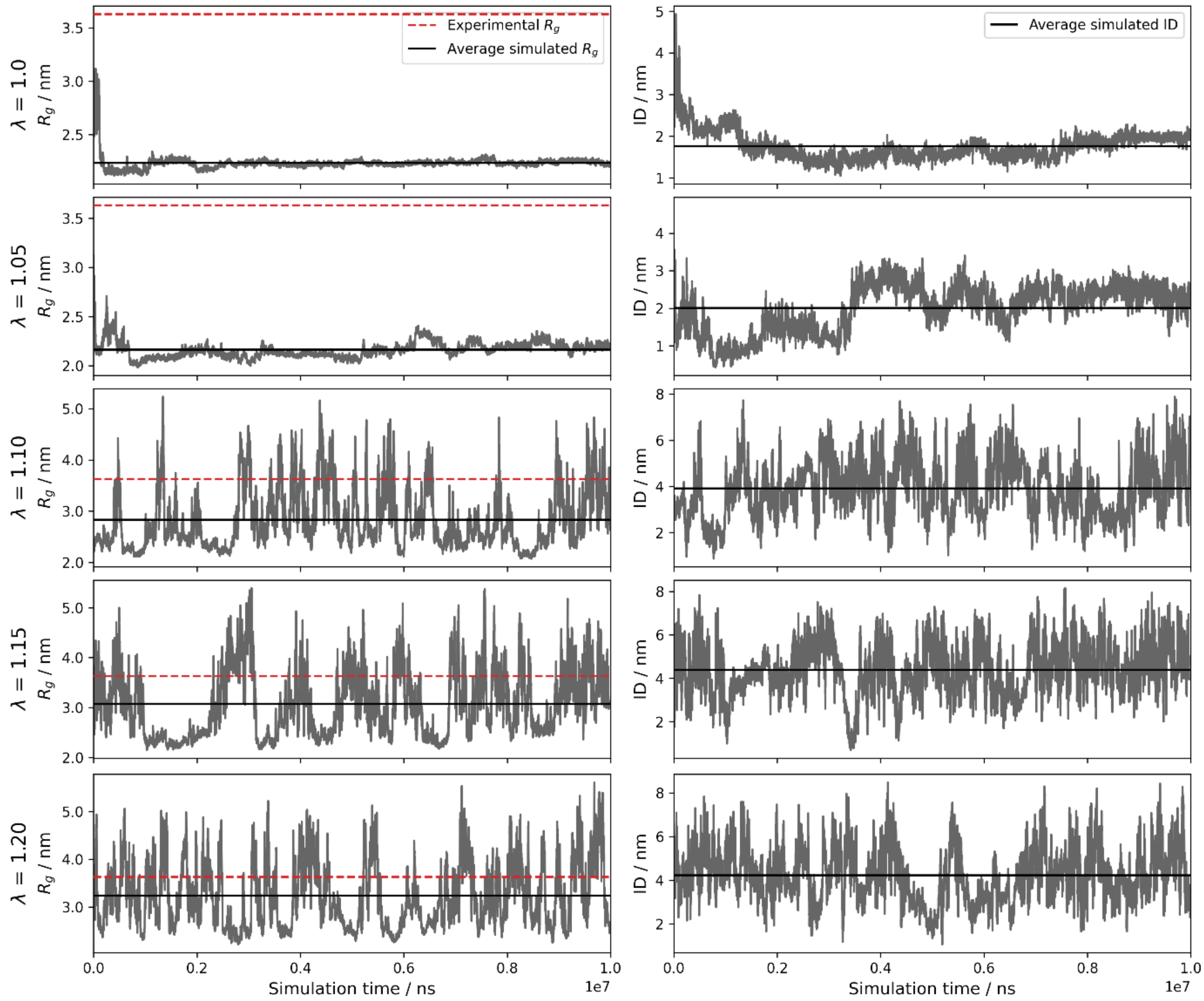


**Figure S4. Simulations of *Sc*LPMO10C** The graphs show time series of the radius of gyration ($R_g$; left-hand panels) and interdomain distance (ID; right-hand panels) for simulations of wild-type *Sc*LPMO10C with different modifications of the strength of protein-water interactions, λ.

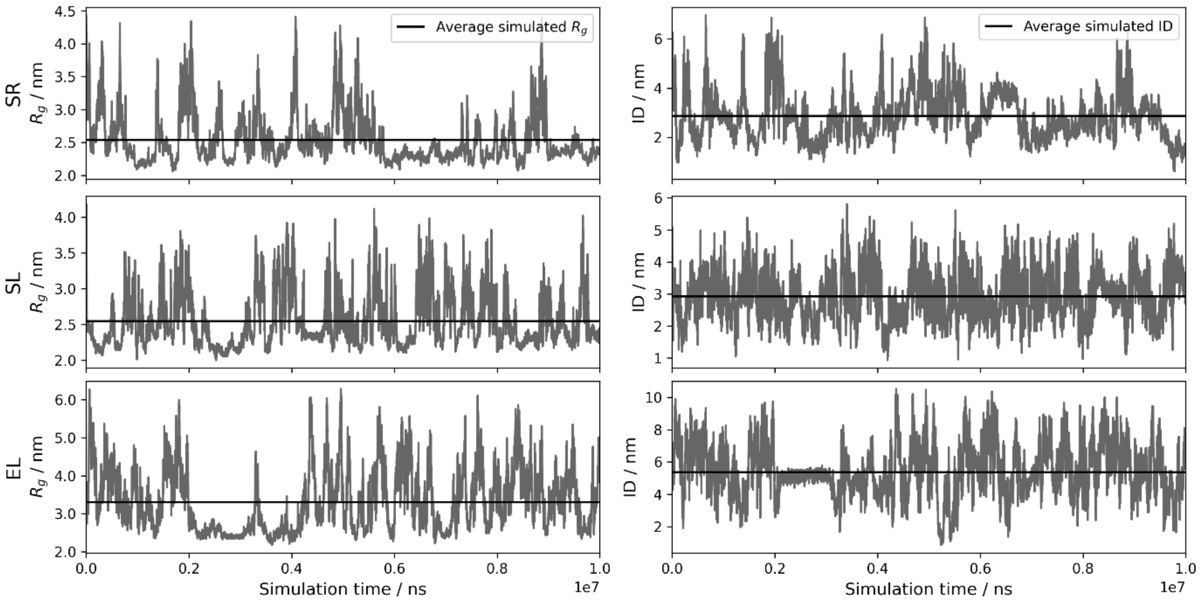


**Figure S5. Simulations of *Sc*LPMO10C linker variants.** The graphs show time series of the radius of gyration ($R_g$; left-hand panels) and interdomain distance (ID; right-hand panels) for simulations of *Sc*LPMO10C linker variants with serine-rich linker (SR), shortened linker (SL), and extended linker (EL).

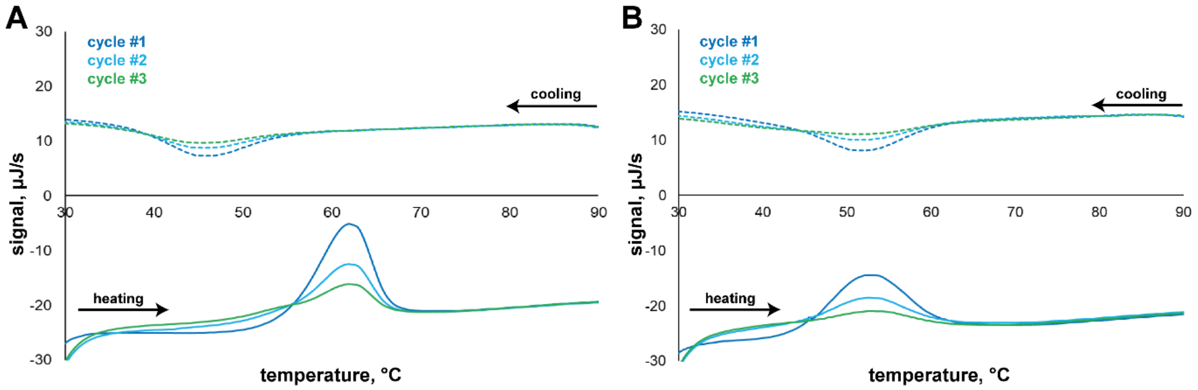


**Figure S6. Unfolding and refolding of the individual domains of *Sc*LPMO10C, *Sc*AA10 (A) and *Sc*CBM2 (B).** The samples subjected to DSC analysis contained 1 g/L protein in sodium phosphate buffer, pH 6.0. One cycle consist of a heating phase (25-90 °C) followed by a cooling phase (90-25 °C) at a rate of 1 °C/min. Note that the melting curves were plotted using raw unprocessed data; the buffer baseline was not subtracted from the protein signal.

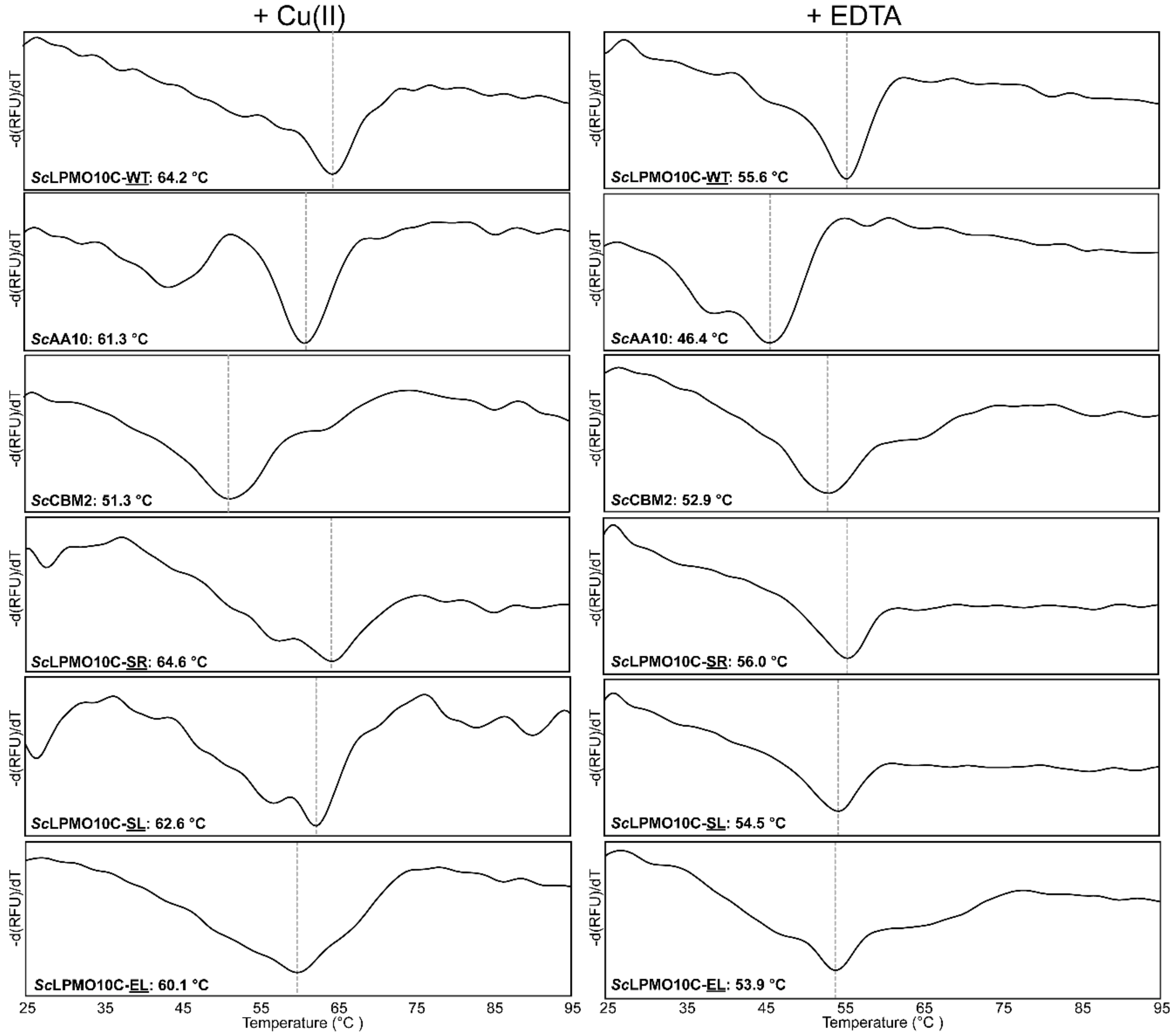


**Figure S7. Thermal stability of *Sc*LPMO10C variants measured by differential scanning fluorimetry (DSF)**. The plots show the melting curves and apparent melting temperatures $T_{m(app)}$ for copper-saturated *Sc*LPMO10C-WT, *Sc*AA10, *Sc*CBM2 and the three linker variants (SR, SL and EL). The derivative of the fluorescence signal (-dRFU/dT, where "RFU" stands for relative fluorescence units") is plotted as a function of the temperature. The reactions contained 0.1 g/L protein in 50 mM sodium phosphate, pH 6.0, and were heated from 25 °C to 97 °C, at a rate of 1.5 °C /min, in the presence of SYPRO orange (a fluorescent dye). The scans were performed four times for each protein and the Figure show a typical scan for each protein. The proteins were copper-saturated as described in Material & Methods prior to the scan (left panels) and 5 mM EDTA was added to the samples shown in the right panels prior to analysis to remove copper from the LPMOs. All apparent melting temperatures $T_{m(app)}$had standard deviations below ± 0.5 °C (n= 4).

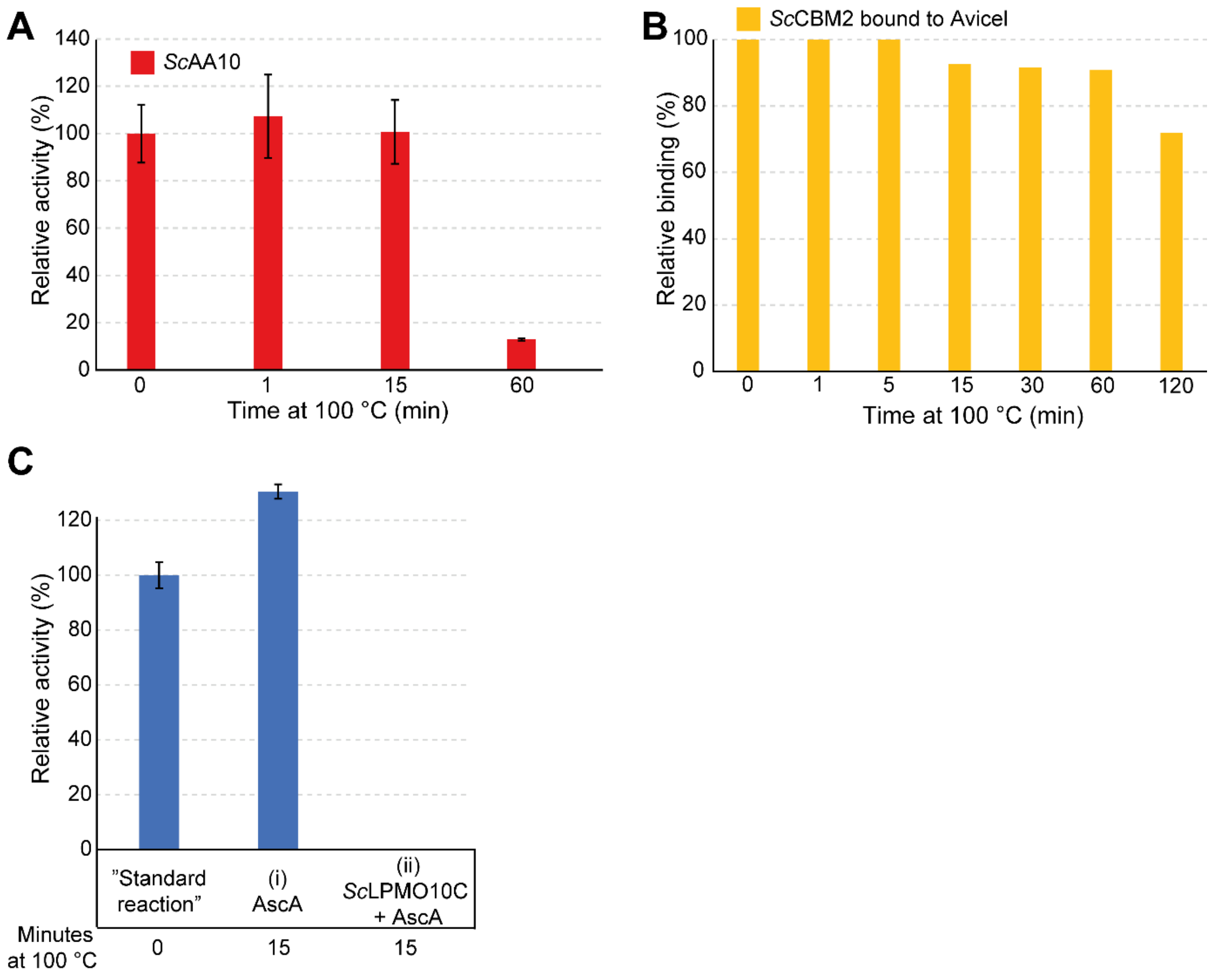


**Figure S8. Residual activity after boiling.** Panel A shows residual activity of *Sc*AA10 exposed to boiling for 0 – 60 minutes. Panel B shows relative binding of the CBM2 to Avicel after pre-exposure of this domain to boiling for 0 – 120 minutes. Panel C shows control reactions in which (i) ascorbic acid (AscA) or (ii) AscA and *Sc*LPMO10C were boiled prior to starting the reactions by adding *Sc*LPMO10C and Avicel to (i) and Avicel to (ii). In the end all reactions contained 10 g/L Avicel, 1 mM AscA, 1 µM *Sc*LPMO10C in 50 mM sodium phosphate buffer, pH 6.0. These reactions were compared to a "standard reaction" that was not exposed to heat containing 1 µM *Sc*LPMO10C-WT, 10 g/L Avicel, and 1 mM AscA in 50 mM sodium phosphate buffer, pH 6.0. All reactions were incubated for 24 h in a thermomixer set to 30 °C and 800 rpm followed by filtration and *Tf*Cel6A degradation of the soluble fraction. The reactions shown in panels A and C were performed in triplicates and the error bars show ±S.D. (n=3).
